# Human 14-3-3 proteins site-selectively bind the mutational hotspot region of SARS-CoV-2 nucleoprotein modulating its phosphoregulation

**DOI:** 10.1101/2021.12.23.474009

**Authors:** Kristina V. Tugaeva, Andrey A. Sysoev, Anna A. Kapitonova, Jake L. R. Smith, Phillip Zhu, Richard B. Cooley, Alfred A. Antson, Nikolai N. Sluchanko

## Abstract

Phosphorylated within its Ser/Arg-rich region, the SARS-CoV-2 nucleoprotein (N) recruits the phosphopeptide-binding human 14-3-3 proteins that play a well-recognized role in replication of many viruses. Here we use genetic code expansion to demonstrate that phosphorylation of SARS-CoV-2 N at either of two pseudo-repeats centered at Ser197 and Thr205 is sufficient for 14-3-3 binding. According to fluorescence anisotropy, the pT205-motif, present in SARS-CoV-2 but not in SARS-CoV, is preferred over the pS197-motif by all seven human 14-3-3 isoforms, which display unforeseen pT205/pS197 binding selectivity hierarchy. Crystal structures demonstrate that pS197 and pT205 are mutually exclusive 14-3-3-binding sites, whereas SAXS and biochemical data indicate 14-3-3 binding occludes the Ser/Arg-rich region, inhibiting its dephosphorylation. This Ser/Arg-rich region of N is highly prone to mutations, as exemplified by the Omicron and Delta variants, with our data suggesting how the strength of its 14-3-3 binding can be linked with the replicative fitness of the virus.

## Introduction

The 46-kDa nucleoprotein (N) is common for single-stranded RNA viruses, including positive-sense coronaviruses, and is responsible for replication, packaging and storage of viral genomic RNA ^1–3^. Being one of the most abundant viral proteins, N can reach up to 1% of the total cellular protein in coronavirus-infected cells ^3–5^ and is a major factor of pathogenicity ^6^. Sharing 89.1% sequence identity with SARS-CoV N, SARS-CoV-2 N consists of the structured N-terminal RNA-binding domain ^7^, the structured C-terminal dimerization domain ^8^, disordered tails and long interdomain linker (residues 176-247) (Fig. 1a) ^9,10^.

**Fig. 1.**
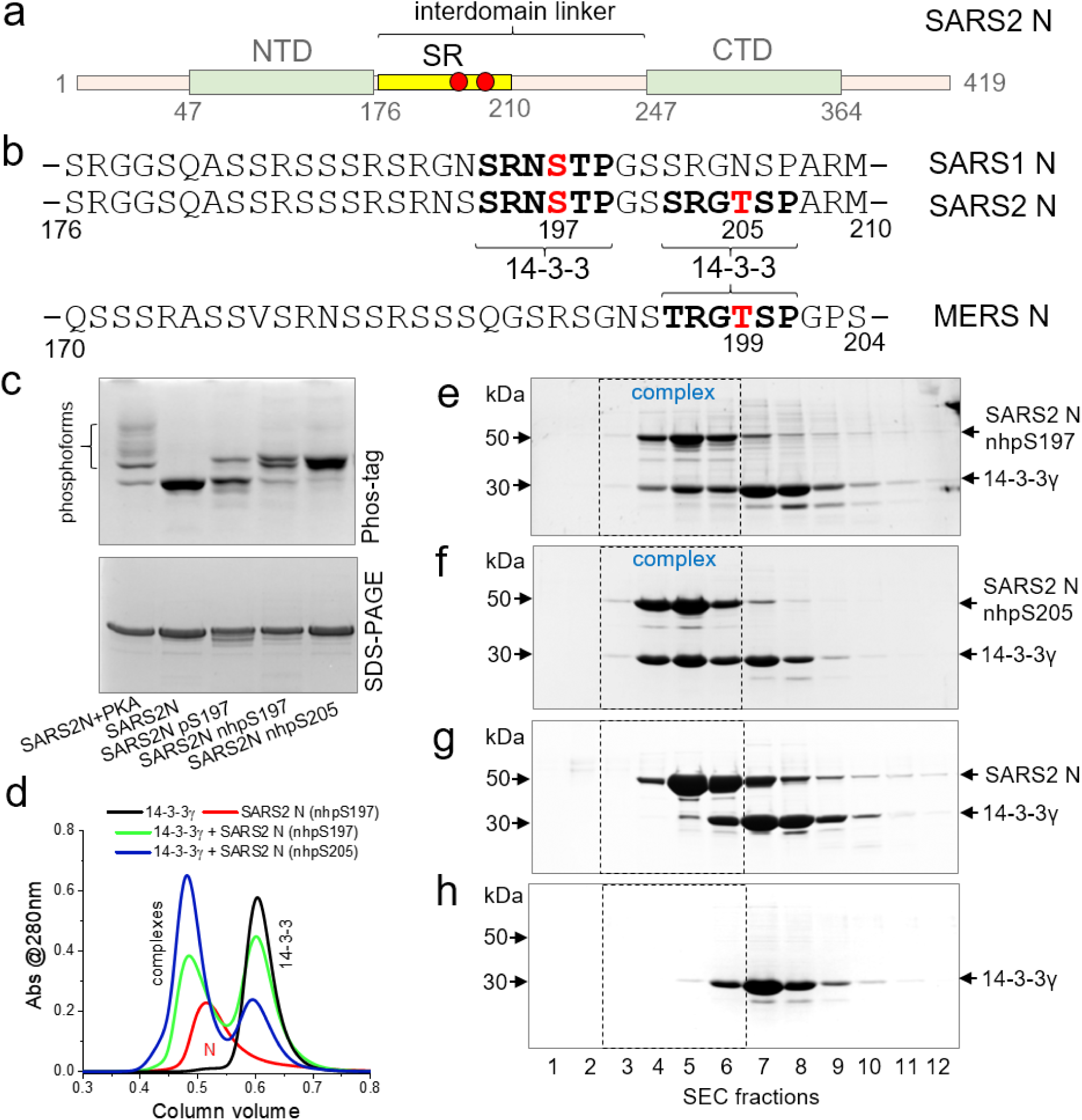
SARS-CoV-2 N recruits human 14-3-3γ using either 197 or 205 phosphosites. a. Schematic representation of N; NTD and CTD, interdomain linker, Ser/Arg-rich region are labeled, Ser197 and Thr205 phosphorylation sites are marked by red circles. b. Sequence alignment of the SR-regions of SARS-CoV-2 and SARS-CoV N proteins, with the phosphosites of interest highlighted in red and potential 14-3-3-binding sites in bold. MERS-CoV N sequence is shown for comparison. Note that SARS-CoV N contains only the first potential 14-3-3-binding site, while MERS-CoV N contains only the second site. c. Phosphorylation-related modification of SARS-CoV-2 N variants analyzed by Phos-tag and SDS-PAGE. d. SEC profiles showing different efficiency of 14-3-3γ complexation with SARS-CoV-2 N containing either nhpS197 or nhpS205. e,f,g,h. Human 14-3-3γ (80 µM) co-elutes with SARS-CoV-2 N (50 µM) containing either nhpS197 (e) or nhpS205 (f) but not with unphosphorylated SARS-CoV-2 N (f) (50 µM) during SEC, as analyzed by SDS-PAGE of the fractions collected. Elution of 14-3-3γ alone (f) is shown for reference. Mass markers (left) and protein positions (right) are indicated. Position of the 14-3-3:N complex on the elution profile is indicated by dashed rectangles. SEC runs were carried out at 220 mM NaCl in the samples and running buffer.

The latter contains a key regulatory Ser/Arg-rich (SR) region (residues ∼176-210) ^11–13^ that mediates higher-order oligomerization of N molecules ^14^, liquid-liquid phase separation with RNA ^11,15^ and is also a hotspot of viral mutations ^15–17^. The SR region of N becomes rapidly phosphorylated by cytoplasmic protein kinases early in infection ^4,18–21^, whereas at its later stages phosphorylation is substantially reduced ^21–23^. Since kinase inhibitors considerably decrease viral titers, phosphorylation of N is critical for the virus fitness ^20,22–24^. Specifically, phosphorylation triggers N binding to 14-3-3 ^25,26^ – the multi-functional phosphopeptide-binding protein-protein interaction hub represented by seven isoforms in human (β, γ, ε, ζ, η, σ, τ) ^27,28^. Upon binding to their specifically phosphorylated protein partners, dimeric 14-3-3 inhibits their dephosphorylation, thereby regulating functional activity, protein-protein interactions and intracellular distribution ^27,29^.

14-3-3 proteins have a widely recognized role in replication of many viruses ^30^. 14-3-3 isoforms are amongst the top 1% of highest-expressed proteins in a number of tissues including the brain, lungs and gastrointestinal system that are particularly susceptible to coronavirus infection ^26^. A detectable up-regulation of some 14-3-3 isoforms upon SARS-CoV-2 infection was reported ^3^. According to immunofluorescence, immunoprecipitation, siRNA silencing and kinase inhibition data, SARS-CoV N undergoes phosphorylation-dependent association with human 14-3-3 in the cytoplasm of the transfected COS-1 cells, which controls nucleocytoplasmic shuttling of N ^25^. Upon silencing of 14-3-3 by siRNA, N accumulated in the nucleus ^25^.

In previous work, we have shown that phosphorylation of the SR region enables SARS-CoV-2 N recognition by human 14-3-3 proteins, leading to a 2:2 complex ^26^. The use of truncated N variants narrowed the binding site to SRNpS^197^TP ^26^, conserved in SARS-CoV and SARS-CoV-2 (Fig. 1a). However, whether other 14-3-3-binding sites exist and the structural basis for 14-3-3/N interactions remained elusive.

Production of specific phosphorylated forms of N is challenged by the dense distribution of phosphorylatable residues (Fig. 1b), resulting in heterogeneously phosphorylated N forms when incubated with kinases ^26,31^. To ensure specific phosphorylation at selected residues, here we utilized the genetic code expansion approach ^32,33^. We identified two phosphorylated pseudo-repeats centered at Ser197 or Thr205 in SARS-CoV-2 N (Fig. 1b) that individually are sufficient to trigger 14-3-3 binding of the full-length SARS-CoV-2 N. Using high-throughput fluorescence anisotropy, we determined the affinities of both phospho-motifs to all seven human isoforms of 14-3-3, which revealed a peculiar trend of 14-3-3 selectivity towards the SARS-CoV-2 N phosphopeptides. Crystal structures of the corresponding 14-3-3 complexes helped rationalize the observed selectivity and, by capturing residues 193-210 of the functional SR region, provided a structural framework for the analysis of 14-3-3 binding to SARS-CoV-2 N variants.

## Results

### Identification of alternative 14-3-3-binding sites within SARS-CoV-2 N

The key 14-3-3-binding site of SARS-CoV-2 N was proposed to center at pSer197, which is conserved also in the SARS-CoV N protein (Fig. 1b) ^26^. To test this hypothesis directly, we first sought to obtain N protein phosphorylated exclusively at the Ser197 position. Since enzymatic treatment always yielded multiply phosphorylated protein (Fig. 1c), we utilized genetic code expansion to site-specifically install phosphoserine at a genetically programmable TAG stop codon ^33^. Although phosphoserine was successfully incorporated to generate SARS-CoV-2 N pS197 protein, confirmed by the discrete upward shift on a Phos-tag gel electrophoresis, we noticed considerable spontaneous dephosphorylation (Fig. 1c). Therefore, for further analysis we produced the SARS-CoV-2 N protein with a non-hydrolyzable analog of phosphoserine, phosphonomethylalanine (nhpS197), inserted in place of the TAG^197^ codon, which caused the desired shift on Phos-tag gel with a negligible amount of unphosphorylated protein (Fig. 1c). This approach was inspired by effective imitation of phosphoserine by nhpSer in previous 14-3-3 binding studies ^32,34^.

On SEC, human 14-3-3γ co-eluted with SARS-CoV-2 N nhpS197, but not with unphosphorylated SARS-CoV-2 N, confirming the formation of the phosphorylation-dependent complex (Fig. 1d, e) and the role of S197 phosphorylation in recruiting 14-3-3. In accordance, the SARS-CoV N protein, which shares the pS197 site with SARS-CoV-2 N (Fig. 1b), similarly associated with 14-3-3γ in a phosphorylation-dependent manner (Supplementary Fig. 1). We then asked if the natural S197L mutation associated with the SARS-CoV-2 N variant identified in 2020 ^35^, which cannot be phosphorylated at this position, would preclude N from interacting with 14-3-3γ. The S197L mutant was co-expressed with PKA for phosphorylation and then purified similar to the wild-type protein. As a control we produced unphosphorylated N.S197L protein, and differences in their phosphorylation status were confirmed by Phos-tag (Supplementary Fig. 2a). Surprisingly, the S197L mutant showed a pronounced phosphorylation-dependent interaction with 14-3-3γ (Supplementary Fig. 2b-f), indicating the presence of alternative 14-3-3-binding sites. The SR region of SARS-CoV-2 N contains a pseudo-repeat around Thr205 (SRG**T**SP), which has a sequence similar to the Ser197 site (SRN**S**TP), undergoes phosphorylation *in vivo* ^4,18^ and by PKA *in vitro* ^26^, and is the best candidate 14-3-3-binding motif.

To test whether phosphorylation at Thr205 was sufficient for N to bind 14-3-3, we expressed full-length SARS-CoV-2 N with nhpSer at this position. Phos-tag gels confirmed near homogenous modification (Fig. 1c) and indeed this phosphosite alone was able to trigger 14-3-3 binding as supported by its co-elution with 14-3-3γ on SEC (Fig. 1f-h). In fact, it bound human 14-3-3γ even more efficiently than the nhpS197 counterpart, i.e. ∼60% vs. 32% of bound 14-3-3γ fraction, respectively, according to the densitometric analysis (Fig. 1d, e, f). Interestingly, while the T205 site is not conserved in SARS-CoV N, the S197 site is (Fig. 1b) ^26^. However, the SR region of MERS N lacks the pS197 equivalent and instead contains a RGT^199^SP motif that is identical to the SARS-CoV-2 N pentapeptide around Thr205 (Fig. 1b). By analogy, it is likely involved in 14-3-3 recognition, although phosphorylation of the MERS N SR region remains to be studied. These considerations indicate that the pS197 and pT205 pseudo-repeats of SARS-CoV-2 N are alternative 14-3-3-binding sites within the regulatory SR region.

**Table 1.**
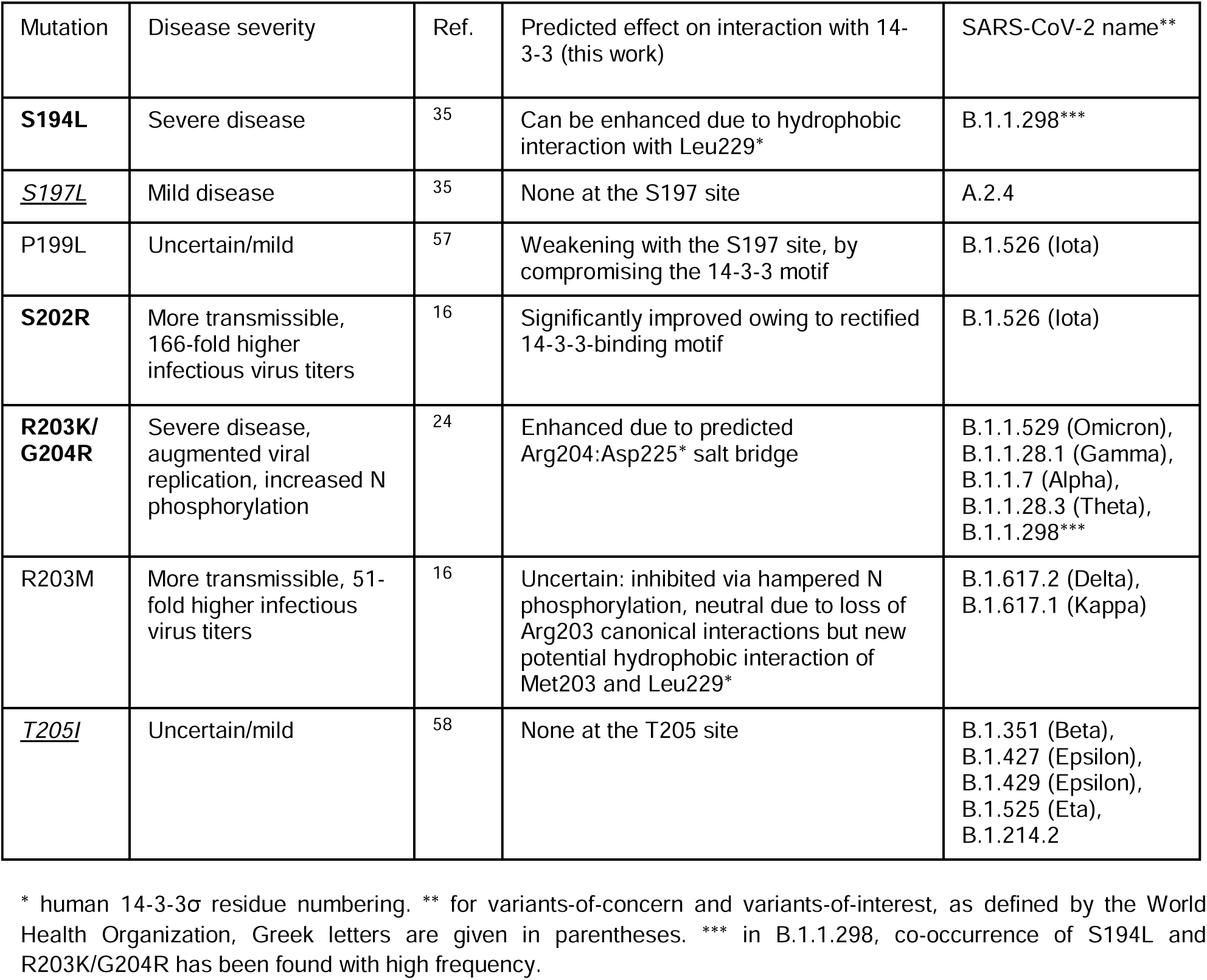
Coupling between the effect of SARS-CoV-2 N mutations and 14-3-3 binding. Bold font indicates prediction of the most pronounced stabilizing effect on 14-3-3 binding. Underlined

### Affinity profiles of Ser197 and Thr205 phosphopeptides towards human 14-3-3 isoforms

Synthetic phosphopeptides SSRNpS^197^TPGSS and SSRGpT^205^SPARM, each bearing an added tryptophan residue for more accurate quantification, were labeled by fluorescein 5-isothiocyanate (FITC) and used to determine binding affinities against all seven human 14-3-3 isoforms by fluorescence anisotropy. For this, homotypic MBP-tagged full-length 14-3-3 constructs were used to obtain highly pure protein in an efficient manner ^28^. SEC-MALS analysis of MBP-14-3-3ζ confirms that the MBP fusion does not disrupt the functional dimeric state of 14-3-3 (Supplementary Fig. 3).

Peptide binding to 14-3-3 was saturable and specific, and the MBP tag did not change the *K*_D_ (Fig. 2a). The same FITC-labeled phosphopeptide failed to bind to noncognate protein lysozyme; likewise, free FITC did not show significant binding to MBP-14-3-3 (Fig. 2a). In addition, the FITC-peptide complexes with 14-3-3 could be disrupted by titration with either the corresponding unlabeled peptides or the canonical phosphopeptide from unrelated protein HSPB6 (pB6) ^36^ (Supplementary Fig. 4). Therefore, the chosen N peptides bind specifically to the common peptide-binding grooves of 14-3-3.

**Fig. 2.**
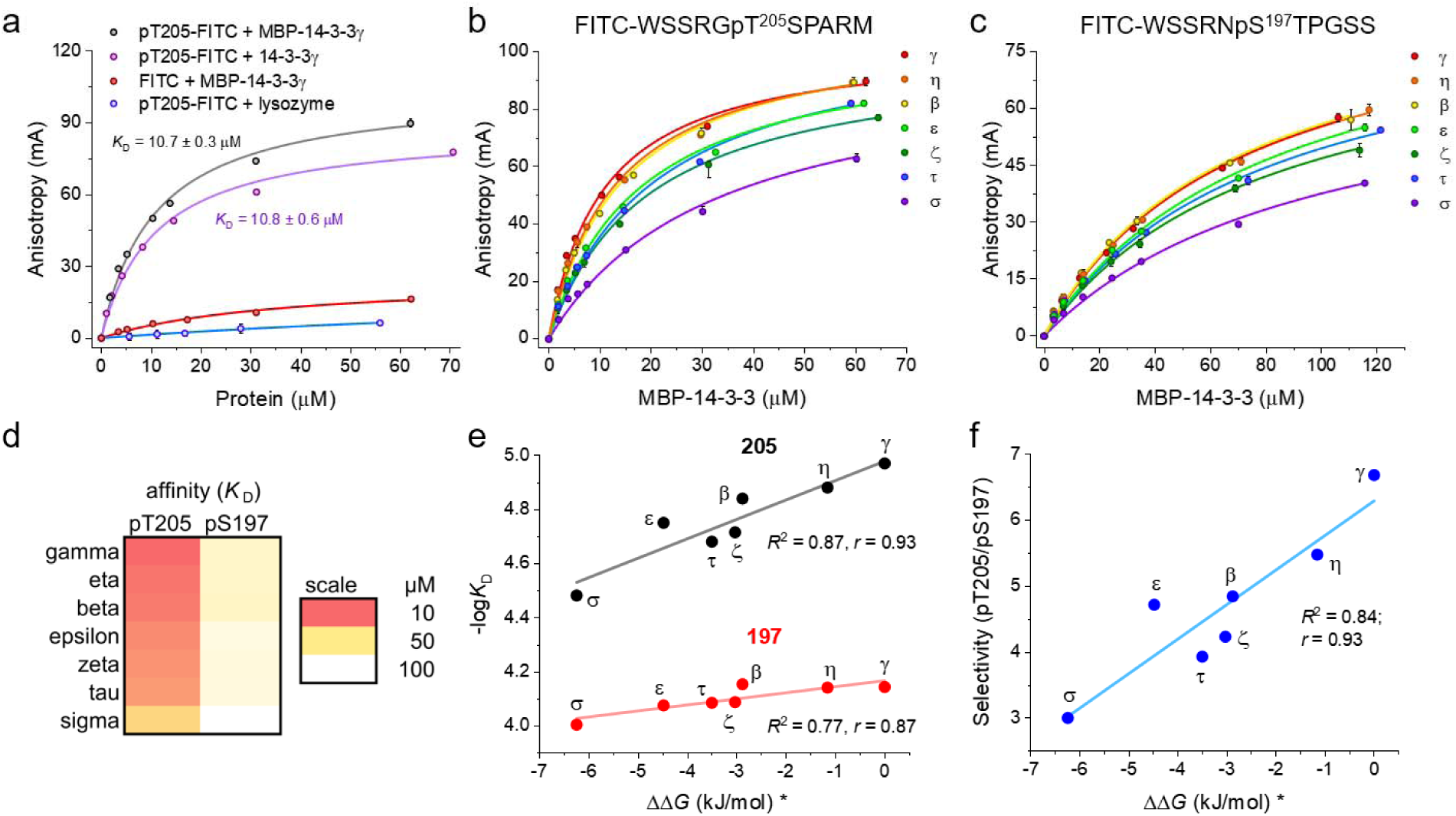
Affinity and selectivity of SARS-CoV-2 N phosphopeptide binding to the human 14-3-3 proteins. a. Fluorescence anisotropy titration data for 14-3-3γ or MBP-14-3-3γ binding to the FITC-labeled SARS-CoV-2 N pT205 peptide. The corresponding apparent *K*_D_ values (mean ± SD, *n*=3) are shown. Titration of the same peptide by lysozyme and titration of free FITC by MBP-14-3-3γ are shown as controls. b, c. Titration of FITC-labeled SARS-CoV-2 N phosphopeptides pT205 (b) or pS197 (c) by the seven human 14-3-3 isoforms. The data (mean ± SD, *n*=3) are color-coded with respect to the 14-3-3 isoforms; fitting curves and the *K*_D_ values at equilibrium are found in Supplementary Fig. 5. In a-c, background anisotropy was subtracted from all values. d. Affinity profiles of the SARS-CoV-2 N phosphopeptides versus 14-3-3 isoforms shown as heatmap gradients according to the *K*_D_ scale on the right. The source *K*_D_ values are found in Supplementary Table 1. e, f. Affinity profiles presented as -log*K*_D_ values (e) or selectivity coefficients (f) are correlated with the proteome-wide affinity hierarchy of the 14-3-3 isoforms represented as average ΔΔ*G* values for the individual isoforms (indicated by asterisks) ^28^. Coefficient of determination *R*^2^ and Pearson’s *r* coefficient characterizing the linear correlations are indicated.

Both SARS-CoV-2 N phosphopeptides showed saturable interaction with all seven human 14-3-3 isoforms, enabling *K*_D_ determination (Fig. 2b and c, Supplementary Fig. 5). The average affinities of 14-3-3 to the SARS-CoV-2 N phosphopeptides were somewhat lower than to the pB6 peptide, yet were in a comparable micromolar range (Supplementary Table 1 and Fig. 5). For each SARS-CoV-2 N peptide, we detected a remarkable peptide-affinity trend following the general peptide-affinity hierarchy of 14-3-3 isoforms reported recently (Fig. 2d and e) ^28^. For the pT205 peptide, the binding affinity to 14-3-3 decreased from gamma (γ) > eta (η) > beta (β) > epsilon (ε) > zeta (ζ) > tau (τ) > sigma (σ); the order for the pS197 peptide was: β ∼ γ ∼ η > ζ ∼ τ ∼ ε > σ (Fig. 2d). As a result, transformation of the *K*_D_ values into -log*K*_D_ showed that both trends correlate well with the interactome-wide affinity hierarchy of the 14-3-3 isoforms (Pearson’s *r*=0.87 for pS197 and *r*=0.93 for pT205, Fig. 2e) ^28^.

The affinity profiles of 14-3-3 for the two SARS-CoV-2 N peptides were significantly shifted and nonparallel to each other. Due to the shift, the affinity of the weakest pT205 peptide binder, 14-3-3σ (*K*_D_ ∼33 μM), was still ∼2 times tighter than for the strongest pS197 binders, 14-3-3β/γ/η (*K*_D_ ∼70 μM). Overall, the 14-3-3 family covered affinities to SARS-CoV-2 phosphopeptides differing by an order of magnitude [∼10-fold *K*_D_ ratio between the weakest (14-3-3σ/pS197) and the strongest (14-3-3γ/pT205) pair], despite the very similar sequences of the peptides and the absolutely conserved 14-3-3 grooves. The average affinity of human 14-3-3 proteins to the studied SARS-CoV-2 phosphopeptides (*K*_D_ ∼50 μM) is roughly in the same order of magnitude as the average affinity of twenty three unrelated peptides analyzed previously (*K*_D_ ∼45 μM) ^28^. This suggests viable competition of the SARS-CoV-2 N phosphopeptides with the host cell phosphopeptides for the 14-3-3 binding, especially with the weaker interacting portions of the interactome.

### Selectivity trend of 14-3-3 isoforms to SARS-CoV-2 N phosphopeptides

Curiously, the affinities to all 14-3-3 isoforms were consistently tighter for the pT205 than for the pS197 peptide (4.3-fold ratio of the average *K*_D_ values) (Supplementary Table 1). Moreover, we found a remarkable selectivity trend reflecting a linear correlation of the *K*_D_ ratio for the 14-3-3 isoforms with their general affinity hierarchy (*R*^2^=0.84, Pearson’s *r*=0.93; Fig. 2f) ^28^. In other words, not only does 14-3-3γ turn out to be the strongest binder, it is also the most selective isoform with respect to the SARS-CoV-2 phosphopeptides pS197 and pT205 (the *K*_D; 197_/*K*_D; 205_ ratio of ∼7), while 14-3-3σ is the weakest and simultaneously the least selective (Fig. 2f). Yet, even for 14-3-3σ the affinity to the pT205 peptide was ∼3 times tighter than for the pS197 peptide. This observation was totally unexpected given the very similar sequences of both peptides, implying that the corresponding protein-peptide complexes feature different interfaces and chemical contacts. To understand the molecular basis of these phenomena, we crystallized 14-3-3σ complexes with both phosphopeptides and determined structures by X-ray crystallography at high resolution (Supplementary Table 2).

### Crystallography explains the selectivity of 14-3-3 for SARS-CoV-2 N phosphopeptides

X-ray structures of 14-3-3σ with both peptides had one 14-3-3 dimer in the asymmetric unit, where each protomer was occupied by the target phosphopeptide. Eight out of ten pS197 phosphopeptide residues and all ten residues for the pT205 phosphopeptide are clearly defined in electron density, revealing key protein/peptide interactions. Residues at +1 and −1 positions with respect to each phosphosite are H-bonded to the side chains of the conserved 14-3-3σ residues Asn175 and Asn226, while hydroxyl groups of both Thr198 and Ser206 of the peptides (position +1) are identically involved in H-bond formation with Asn175 and the conserved Lys122 residue of 14-3-3σ (Fig. 3a, b and Supplementary Fig. 6). An additional H-bond in the case of the pS197 peptide between the side chain of its Asn196 residue and Asn226 of 14-3-3σ is replaced by the additional hydrophobic contact in the case of the pT205 peptide, formed between the methyl group of phospho-Thr205 residue and the side chain of 14-3-3σ’s Val178. In contrast to the nearly equivalent central six-residue segments of both peptides (Cα RMSD=0.7 Å), crystal structures revealed significant differences in their C-terminal regions. In the case of the pS197 peptide, we could not detect any stabilizing *in-trans* interactions after Pro199. The lack of defined electron density for the last two Ser residues of this peptide indicate flexibility and hence inability to form stabilizing interactions (Fig. 3a). On the contrary, the pT205 peptide was additionally stabilized *in-trans* by i) the hydrophobic contacts of its Ala208 with Val46 of 14-3-3 and Met210 with Phe118 and Ile168 of 14-3-3, ii) the H-bonding interaction between the peptide’s carbonyl of Ala208 with the side chain of Ser45 of 14-3-3σ and iii) the salt bridge between the peptide’s Arg209 with Glu14 of 14-3-3σ (Fig. 3b). Interestingly, Glu14 is absolutely conserved in all human isoforms of 14-3-3 (Supplementary Fig. 7). Therefore, the salt bridge interaction involving Arg209 is likely shared by all 14-3-3 isoforms and, together with factors i)-ii), confer the consistently higher affinity of the pT205 peptide versus pS197 peptide to 14-3-3 proteins, as observed biochemically (Fig. 1 and 2).

**Fig. 3.**
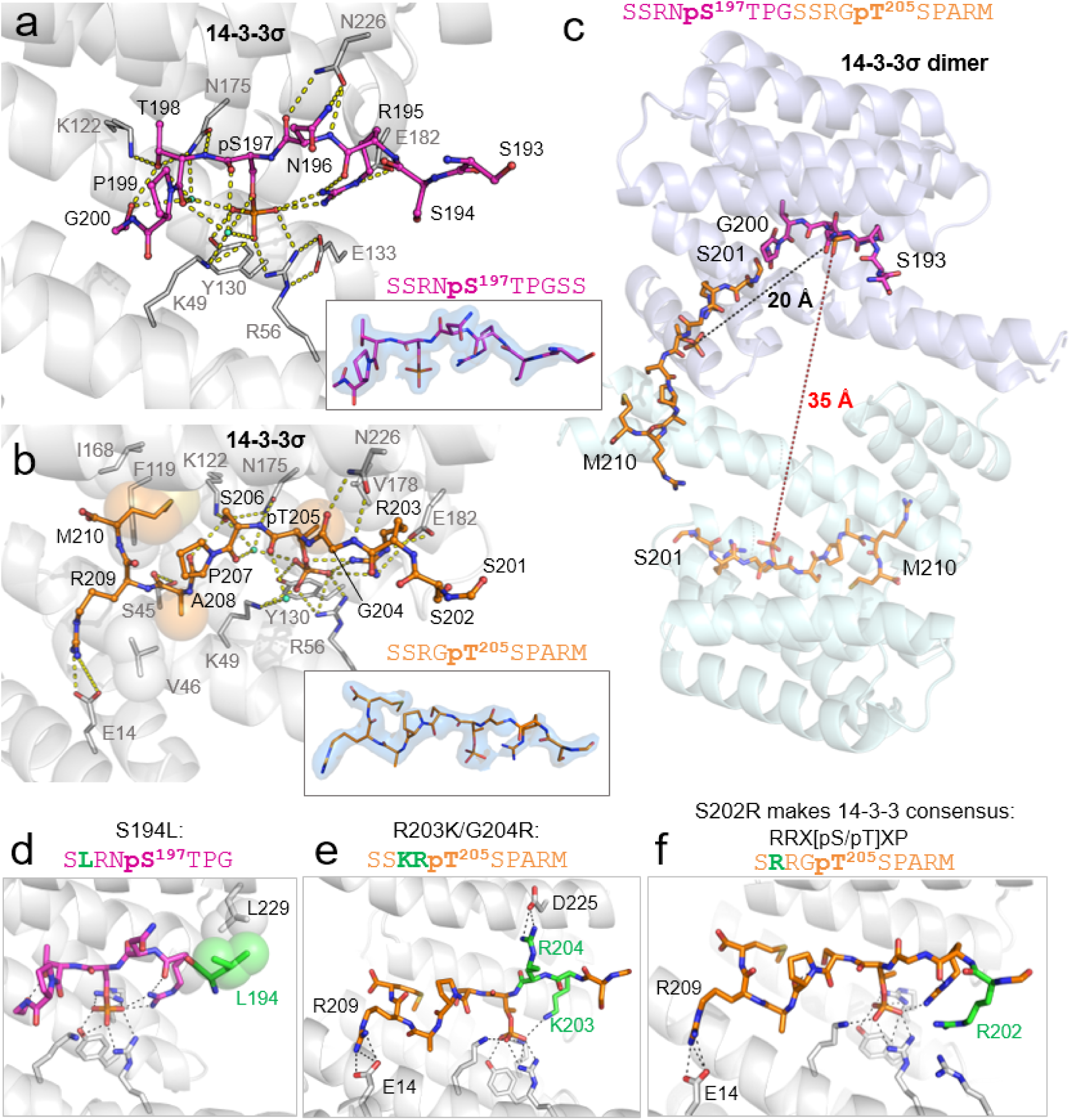
Structural basis for the 14-3-3σ selectivity towards the two alternative phosphosites within SARS-CoV-2 N. a,b. Molecular contacts between 14-3-3σ (gray cartoon) and the pS197 phosphopeptide (magenta sticks) (a) or the pT205 phosphopeptide (orange sticks) (b). Hydrogen bonds are indicated by yellow dashed lines, hydrophobic interactions are indicated by large semi-transparent spheres, water molecules mediating 14-3-3-peptide contacts are in aquamarine. 2Fo-Fc electron density maps for the phosphopeptides contoured at 1σ are shown as semi-transparent blue surfaces in the inserts. Sequences of the phosphopeptides are shown above the inserts. c. A model showing that the two phosphopeptides (magenta and orange sticks) of the same N chain cannot occupy both 14-3-3σ subunits (shown as semi-transparent cartoon) simultaneously. The pT205 peptide is depicted either bound in the bottom 14-3-3 groove (semi-transparent sticks) or as a continuation of the pS197 peptide bound in the top 14-3-3 groove. Distances between two phospho-groups in one continuous peptide (black dashed line) or as separate 14-3-3-bound entities (red dashed line) are indicated. d-f. SARS-CoV-2 N mutations improving fitness of the virus potentially affect 14-3-3 binding as illustrated by structural models showing the expected interaction of the mutated N peptides with 14-3-3 based on our crystal structures. The mutated residues are shown in green, polar contacts are shown by black dashed lines, hydrophobic interactions are indicated by semi-transparent spheres.

These peptides of the same SARS-CoV-2 N polypeptide chain cannot simultaneously occupy both grooves of the 14-3-3 dimer, as the pS197 and pT205 phosphosites are only 7 residues apart (the inter-phosphate distance is 20 Å). For bidentate binding to 14-3-3, at least 15 extended residues ^37^ are needed to sufficiently separate the phosphosites (e.g. 22 in the 5N6N structure ^38^) and cover the required ∼35 Å distance between the two binding sites of the 14-3-3 dimer (Fig. 3c). While a mixed pS197/pT205 binding within the 2:2 SARS-CoV-2 N/14-3-3 complex cannot be excluded, binding of two pT205 motifs to the 14-3-3 dimer would be thermodynamically preferred, unless mutations in the SR region change this selectivity. Such bidentate 14-3-3 dimer binding using either of these two phosphosites is expected to increase the affinity in the context of the full-length proteins ∼10-fold ^26^.

### Structure-based prediction of effects of the N mutations on 14-3-3 binding

The conformation of the 14-3-3-bound peptides allowed us to rationalize the effect of N mutations present in different coronavirus variants (Table 1) by predicting their influence on 14-3-3 binding. First, due to the phosphorylation-dependent nature of 14-3-3 binding, the S197L and T205I mutations of N simply disqualify the corresponding site. Interestingly, the S197L mutation has a statistically mild outcome ^35^. The T205I mutation, present in multiple variants of SARS-CoV-2, has either mild outcome or co-occurs with multiple mutations in other SARS-CoV-2 genes (https://viralzone.expasy.org/9556), making predictions about its physiological effect challenging. The P199L mutation can decrease the affinity to 14-3-3 at the pS197 site because this position is canonically occupied by Pro ^39^, but does not compromise 14-3-3 binding at the pT205 site. The S194L mutation, associated with severe COVID-19 ^35^, can site-specifically strengthen 14-3-3 binding due to the additional hydrophobic contacts between Leu194 and a leucine residue of 14-3-3 facing this position in the crystal structure (Fig. 3d). The widespread combined R203K/G204R mutation present in at least several variants-of-concern (e.g., Alpha, Gamma, Omicron) is expected to enhance phosphorylation of N ^24^ and affect its complex with 14-3-3. While the R203K mutation is expected to slightly decrease the *in-cis* contact with the phosphate group, the Arg204 side chain can reach the conserved Asp225 of 14-3-3 to make a salt bridge (Fig. 3e). The S202R mutation has the most profound effect on the efficiency of N to package RNA and produce new viral particles ^16^, with at least two predictable effects on the 14-3-3 level. First, the S202R mutation can enhance phosphorylation of Thr205, at least by basophilic protein kinases (by forming a canonical sequence RRXT instead of just RXT ^40^). Second, it converts the **S**RGpTSP 14-3-3-binding site from suboptimal to canonical (**R**RGpTSP) ^39^, which would strongly promote 14-3-3 binding (Fig. 3f). The effect of the R203M mutation associated with the Delta variant is difficult to predict at the level of 14-3-3 binding, because this mutation would interfere with Thr205 phosphorylation in the S**M**GT^205^SP context, at least by basophilic protein kinases.

### 14-3-3 occludes the SR region of N and inhibits its dephosphorylation

Unstructured regions of SARS-CoV-2 N restrict its ability to crystallize, therefore we used small-angle X-ray scattering (SAXS) to gain insight into the spatial organization of N in complex with 14-3-3. To make sure that only particles corresponding to the complex were analyzed, we used SEC-SAXS to separate a mixture of phosphorylated N with an excess of 14-3-3γ and ensure scattering data were collected on the intact complex only. This setup also enabled to control the aggregation state of the system, especially important given the inherent higher-order self-association propensity of N ^9,14^. Independent MALS analysis assisted in validating the 2:2 complex stoichiometry (150.8 kDa; expected *M*_w_ is 148.3 kDa) and in selecting the representative region for averaging SAXS curves for the complex (Fig. 4a).

Complex particles had average *R*_g_ of ∼5.5 nm and maximal dimensions of ∼20 nm (Fig. 4b,Supplementary Table 3). Kratky plots confirmed the presence of folded bodies organized in a non-spherical shape, connected by flexible linkers (Fig. 4c). To model this complex, we used available atomistic models of dimeric 14-3-3γ as well as of the NTD, dimeric CTD and the short stable α1-helix in the interdomain linker of SARS-CoV-2 N ^41^. Flexibility of IDRs has been confirmed in different N constructs by NMR ^41^. We used the N S197L construct because it featured minimum extra residues and contained the major 14-3-3-anchoring pT205 motif in both N chains. While keeping the crystallographic 14-3-3/pT205 peptide interface, we used CORAL ^42^ to determine probable positions of the NTD, CTD and α1 parts of N relative to the 14-3-3γ dimer (Fig. 4d). Twenty independently calculated models provided excellent fits to the SAXS data (χ^2^=1.04-1.28) (Fig. 4e), the fits by individual 14-3-3 or N dimers were incomparably worse (χ^2^>200 and 4.5, respectively). While differing by positions of N foldons with respect to the 14-3-3/pT205 core, the complex models revealed that 14-3-3 grooves mask a significant portion (20-25 residues) of the SR N region (Fig. 4d). An analogous, pS197-mediated anchoring of 14-3-3 by the wild-type N protein would occlude the ∼186-211 SR segment, covering an even larger number of phosphorylatable residues (S186, S187, S188, S190, S193, S194, S197, T198, S201, S202, T205, S206 of SARS-CoV-2 N ^26^ or equivalent phosphosites of SARS-CoV N (Fig. 1b)). Since this can profoundly interfere with phosphoregulation of N, we checked this hypothesis by analyzing the effect of 14-3-3γ on N dephosphorylation kinetics *in vitro*. In this experiment, we used phosphorylated SARS-CoV N because co-expression with PKA provided the best experimental window, i.e. the largest electrophoretic mobility shift on Phos-tag gels (Supplementary Fig. 1). Using 25 µM of full-length phosphorylated SARS-CoV N and 60 µM of 14-3-3γ and assuming the *K*_D_ of the 2:2 complex of ∼7 µM (i.e., ∼10 times tighter than for the 14-3-3γ/pSer197 peptide binding), we expected a ∼85% complex formation and therefore a measurable effect of 14-3-3γ on susceptibility of N to dephosphorylation. The presence of human 14-3-3γ has indeed slowed down the phosphatase reaction by >2-fold, yielding a considerable inhibition of SARS-CoV N dephosphorylation in the course of 4 h phosphatase treatment (Fig. 4f, g). Incomplete inhibition is likely explained by the presence of N phosphosites beyond the 14-3-3-protected region and by the dynamic nature of the 14-3-3/pN complex, indicating that 14-3-3 binding modulates the phosphoregulation of N.

**Fig. 4.**
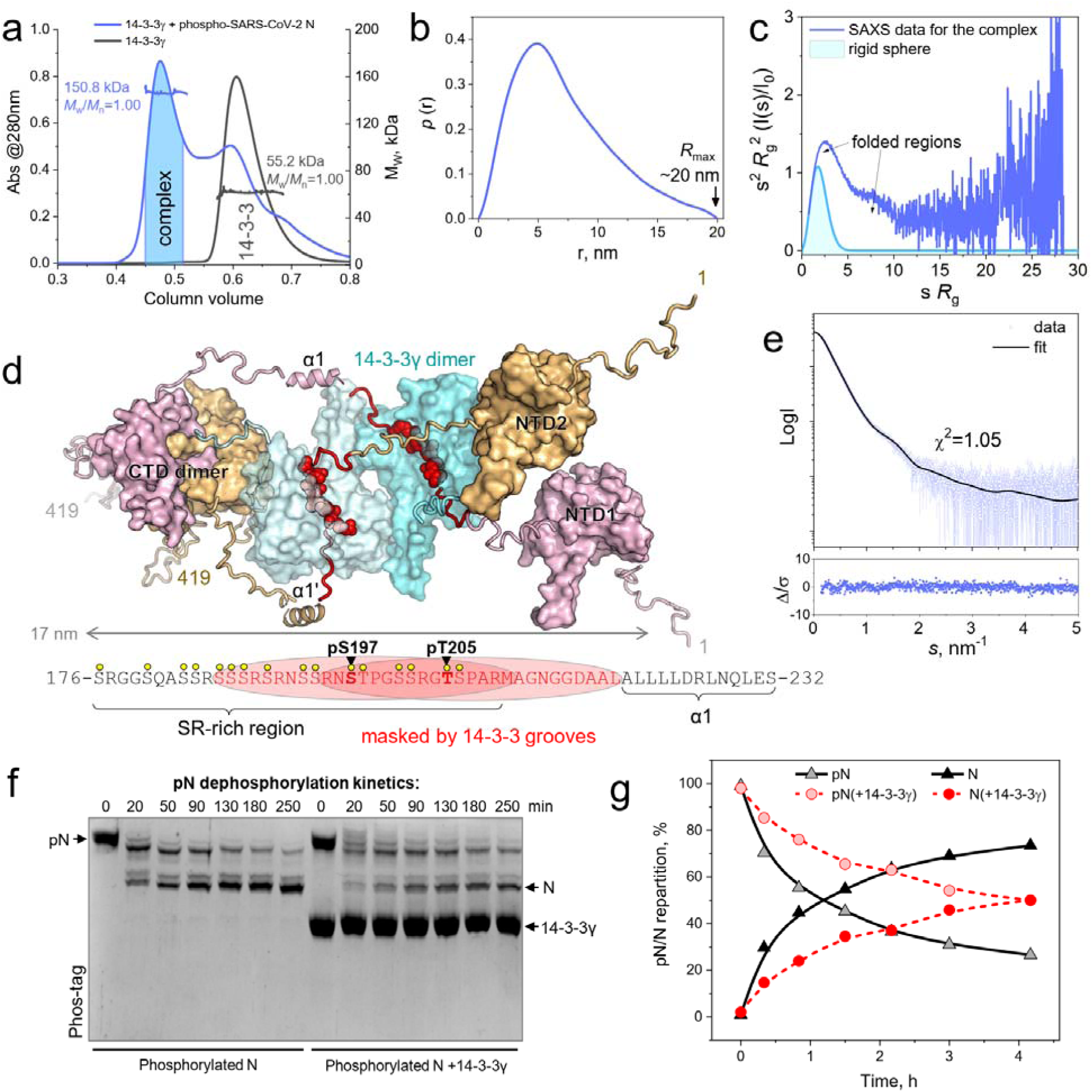
In complex with phosphorylated N, 14-3-3γ occludes part of the SR region and inhibits N dephosphorylation. a. SEC-MALS profiles of 14-3-3γ or its mixture with the phospho-S197L mutant. Average *M*_w_ values and polydispersity indexes (*M*_w_/*M*_n_) for the peaks are indicated. Highlighted is the region used for structural analysis of the complex based on online SAXS. b. Pairwise distance distribution function for the complex. c. Dimensionless Kratky plot for the complex, compared with that for a rigid sphere. The bell-shaped peak and a shoulder corresponding to folded parts of the complex are indicated by arrows. d. One of the possible conformations of the 2:2 assembly based on CORAL modeling ^42^. SARS-CoV-2 N polypeptides are shown by pink and wheat colors, 14-3-3γ dimer is shown by tints of cyan, portions of the interdomain linker of N masked by 14-3-3γ grooves are shown by red, including the 14-3-3γ-bound pT205 phospho-motifs (red spheres in the model) matching those in our 7QIP structure. The foldons of N are named. The regions of N masked by either pS197- or pT205-bound 14-3-3γ are highlighted by red font and ovals on the scheme below. Small yellow circles indicate phosphorylatable residues. e. The fit of the CORAL model presented on **d** to the SAXS data with the associated residuals (Δ/σ). f. Kinetics of dephosphorylation of phosphorylated SARS-CoV N by shrimp alkaline phosphatase in the absence or presence of human 14-3-3γ, analyzed by Phos-tag SDS-PAGE. Positions of proteins are indicated by arrows. g. Densitometric analysis of the gel in f reveals the effect of 14-3-3γ binding on repartition of SARS-CoV N between the phosphorylated (pN) and dephosphorylated (N) forms.

## Discussion

Unlike mutations in S protein, that can often be rationalized via their direct effect on viral entry, the effects of mutations in N remain largely unknown. Curiously, the majority of the widespread N mutations beneficial for the virus are mapped to a very small segment, residues 194-205 (Fig. 1a,Table 1) ^16,17,43^. In fact, all variants-of-concern defined by the World Health Organization feature at least one mutation in the 199-205 subregion ^16^, which coincides with the maximum Shannon entropy in the SARS-CoV-2 genome centered at these positions ^17^. Unfortunately, this mutational hotspot SR region of N is disordered, which reduces our ability to deconvolute the underlying effects of relevant mutations at the structural and functional level.

Recent work using the viral-like particles to assess evolved SARS-CoV-2 N variants has shown that the S202R and R203M mutations vastly improve RNA packaging mechanisms and production of higher titers of infectious virions (166- and 51-fold, respectively) ^16^. Since the improved replicative fitness of the virus correlates well with the increased levels of N phosphorylation ^20,24^, factors promoting N phosphorylation or postponing its dephosphorylation appear particularly relevant for the success of the virus. Noteworthily, 14-3-3 proteins play a well-recognized role in both the replication of many viruses ^30^ and in protecting their bound phospho-clients from dephosphorylation ^44,45^, which is likely the case for phosphorylated SARS-CoV-2 N when bound to 14-3-3.

We have shown that multiply phosphorylated SARS-CoV-2 N bearing several physiologically-relevant phosphosites ^4,18^ can be successfully reconstituted *in vitro* to induce its interaction with 14-3-3 ^26^. Our present study identified that 14-3-3 site-selectively recognize two adjacent phosphorylated pseudo-repeats centered at Ser197 and Thr205 of SARS-CoV-2 N. Importantly, this was done in the context of full-length proteins and could only be achieved using the latest genetic code expansion approaches enabling site-specific incorporation of phosphoserine and a non-hydrolyzable analog thereof ^32,33^. Switching to the protein-peptide binding format, we used fluorescence anisotropy to show that both SARS-CoV-2 N phosphopeptides display the characteristic peptide-affinity profiles to the human 14-3-3 isoforms ^28^. Unexpectedly, the pT205 fragment interacted with all seven 14-3-3 protein isoforms with a consistently tighter affinity than the pS197 fragment. Moreover, the interaction of the pT205 peptide with the weakest isoform, 14-3-3σ, was still stronger than the interaction of the pS197 peptide with the strongest isoform, 14-3-3γ (Fig. 2).

Most importantly, we identified the remarkable selectivity profile for the 14-3-3 family, a phenomenon that, to the best of our knowledge, has not been previously recognized. This selectivity trend correlated well (*R*^2^=0.84; *r*=0.93) with the general peptide-affinity hierarchy of the 14-3-3 isoforms ^28^. Accordingly, not only the strongest peptide binder, 14-3-3γ, binds the pT205 peptide stronger, but its selectivity to this peptide over the pS197 peptide is more pronounced than in the case of generally the weakest isoform, 14-3-3σ (Fig. 2f).

To explain the established binding preferences of 14-3-3 proteins toward one of the apparently similar phosphopeptides, we determined crystal structures of 14-3-3σ with both peptides (Fig. 3). The central six-residue portions of the peptides indeed form roughly equivalent contacts with 14-3-3. By contrast, specific stabilizing interactions beyond the central region were observed only in the case of the pT205 peptide. The pS197 and pT205 fragments are mutually exclusive, because their simultaneous placement within the 14-3-3 dimer is sterically constrained (Fig. 3c). Our model of the 14-3-3 interaction with N based on SAXS data strongly indicated that the complex formation occludes regulatory SR region of N, i.e. ∼20-25 residues surrounding each 14-3-3-binding phosphosite, either 197 or 205 (Fig. 4a-e). Given that >10 potential phosphosites are affected, this would result in measurable inhibition of N dephosphorylation by 14-3-3. We’ve tested this hypothesis showing that dephosphorylation of phosphorylated SARS-CoV N is indeed inhibited in the presence of 14-3-3γ (Fig. 4f,g). This is in line with previous observations where 14-3-3 protected other phospho-client proteins from dephosphorylation ^44,45^ and suggests that 14-3-3 binding modulates the phosphoregulation of N.

SARS-CoV-2 N fragments 193-200 and 201-210 co-crystallized with 14-3-3 provide a valuable structural framework for analyzing the effect of the widespread mutations and testing new hypotheses. Collectively, our analyses based on structural, interactomic and bioinformatic data support the hypothesis ^17^ that N mutations improving SARS-CoV-2 fitness positively affect its binding to 14-3-3 (Fig. 3). Affecting N dephosphorylation, 14-3-3 binding can promote N functions relying on phosphorylation ^20–23^. The SR region of N masked by 14-3-3 is flanked by regions 179-184 and 217-222 that appear to drive amyloid aggregation of N in the course of liquid-liquid phase separation in the presence of RNA ^46^. Because phase separation is important for viral replication and is affected by N phosphorylation ^15,21^, phosphorylation-mediated 14-3-3 binding can act as another layer of regulation that is targeted by the widespread mutations. While this warrants further investigation, we note that 14-3-3 binding, mediated by structurally disordered segments containing phosphorylation sites, emerges as a common feature in virus-host interactions ^25,26,28,30,47^.

## Methods

### Peptides and chemicals

SARS-CoV-2 N peptides phosphorylated at Ser197 (WSSRNpSTPGSS) or Thr205 (WSSRGpTSPARM) were synthesized by Severn Biotech (UK). The phosphopeptide of human HSPB6 (pB6; WLRRApSAPLPGLK) was obtained as previously described ^36^. Expected and experimentally determined masses of the peptides are listed in Supplementary Table 4. FITC was from Sigma-Aldrich (cat no. F7250). Phos-tag acrylamide was purchased from Wako Chemicals (AAL-107). Lysozyme from hen egg white was from Roche (cat no. 10837059001). Shrimp alkaline phosphatase (#EF0511, Fermentas) was kindly provided by Dr. Michael Agaphonov. All common chemicals were of the highest purity and quality available, and all solutions were made with milliQ grade water.

### FITC labeling of the SARS-CoV-2 N phosphopeptides

SARS-CoV-2 N peptide labeling was achieved in 0.1 M carbonate buffer, pH 9.0, by mixing a 5 mg/ml solution of the peptide with 1/10 volume of the FITC stock solution (9 mg/ml) in DMSO. The pB6 phosphopeptide ^36^ was labeled according to the same scheme and was used for controls. The mixtures were wrapped with aluminum foil and incubated overnight at room temperature, then the reaction was quenched by the addition of Tris to a final concentration of 1 M. The FITC-labeled peptides were separated from the unreacted label by SEC on a 2 ml Sephadex G10 column equilibrated and run using a 20 mM Tris-HCl buffer, pH 7.5, 150 mM NaCl at a 0.5 ml/min flow rate. The runs were operated by the Prostar 335 system (Varian Inc., Australia) following the full-spectrum absorbance. The resulting spectrum, and absorbance at 280 (*A*_280_) and 493 nm (*A*_493_) in particular, was used to estimate the phosphopeptide concentration (*C*_peptide_), taking into account FITC absorbance at 280 nm using the formula ^48^:

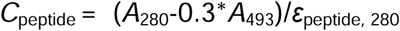

where ε_peptide, 280_ is the extinction coefficient of the peptide at 280 nm (for all peptides in the study 5500 M^-1^ cm^-1^). FITC concentration was determined from the FITC-specific absorbance at 493 nm using the extinction coefficient of 68,000 M^-1^ cm^-1^. The completion of the labeling reaction was confirmed by LC-MS on a Bruker Impact II mass-spectrometer (Supplementary Table 4 and Supplementary Fig. 8). The labeling efficiency was 74-76% for all phosphopeptides. Labeled peptides were stored in aliquots at −80 °C.

### Plasmids and mutagenesis

The SARS-CoV-2 N sequence (Uniprot ID P0DTC9) was cloned as described previously ^26^. The protein contained an N-terminal His_6_-tag cleavable by rhinovirus 3C protease, which left extra residues GPA on the N terminus. The wild-type SARS-CoV-2 N plasmid was used as template to PCR amplify the S197L mutant using the S197L reverse mutagenic primer (5’-CTGGAGT<u>TAA</u>ATTTCTTGAACTG-3’, the mutated codon underlined). For site-specific translational incorporation of phosphoserine or its non-hydrolyzable analog phosphono-methyl-alanine (nhpSer) at positions 197 or 205 of SARS-CoV-2 N using amber codon suppression in autonomous *E. coli* systems ^32,33^, we created the 197TAG and 205TAG mutants using the reverse mutagenic primers (5’-CTGGAGT<u>CTA</u>ATTTCTTGAACTG-3’ for 197TAG or 5’-CAGGAGA<u>CTA</u>TCCCCTACTGCTG-3’ for 205TAG, the mutated codons are underlined) and DNA template being the SARS-CoV-2 N plasmid with a C-terminal His_6_-tag cleavable by 3C protease. The resulting PCR products including the C-terminal cleavable His_6_-tag were treated with NdeI/HindIII restriction endonucleases and then cloned into a pRBC_14-3-3γ/His_6_SUMO_mBAD_43-204_ vector ^33^ pretreated with the same endonucleases. The C-terminal His_6_-tag ensured facile removal of truncated and untagged 1-197 and 1-204 products forming due to the presence in the *E*.*coli* cells of the Release Factor 1 responsible for translation termination at TAG codons ^49^. After cleavage by 3C protease, 197TAG or 205TAG proteins retained extra residues GLEVLFQ on their C terminus. The correctness of all constructs was verified by DNA sequencing in Evrogen, Moscow (see Supplementary Fig. 9). The SARS-CoV N-containing pGEX plasmid (ampicillin resistance) was kindly provided by Prof. Fulvio Reggiori (University of Groningen, The Netherlands).

Untagged full-length human 14-3-3γ (Uniprot ID P61981) cloned into a pET21 vector (ampicillin resistance), the mutants of the C-terminally truncated human 14-3-3σ (residues 1-231), i.e. Clu1 (^159^KKE^161^ → ^159^AAA^161^) and Clu3 (^75^EEK^77^ → ^75^AAA^77^), each containing an N-terminal 3C protease cleavable His_6_-tag and cloned into the pET28 vector (kanamycin resistance) have been described previously ^36^. Homotypic constructs of the full-length human 14-3-3 protein isoforms beta (β), gamma (γ), eta (η), tau (τ), zeta (ζ), epsilon (ε), and sigma (σ), fused to the C terminus of the maltose-binding protein (MBP) ^28^, were kindly shared by the laboratory of Prof. Gilles Travé (Université de Strasbourg, Illkirch, France). Parameters of proteins used are provided in Supplementary Table 5.

### Protein expression and purification

Proteins were expressed in *E*.*coli* BL21(DE3) by the addition of IPTG up to 0.5 mM for 20 h at 25 °C (N proteins), or for 4-6 h at 37 °C (14-3-3 proteins). To ensure phosphorylation of SARS-CoV-2 N, its S197L mutant, or SARS-CoV N in *E*.*coli* cells, target proteins were co-expressed with the catalytically active subunit of mouse protein kinase A (PKA) encoded in a separate pACYC-PKA plasmid (chloramphenicol resistance), essentially as described earlier ^26^. The site-specifically phosphorylated SARS-CoV-2 N protein pS197 was expressed in BL21(DE3) Δ*serB* cells tailored to co-translational phosphoserine incorporation ^49^. This yielded partially phosphorylated purified protein, therefore we also produced the nhpSer197 protein based on the same plasmid containing a TAG^197^ codon (ampicillin resistance) in BL21(DE3) Δ*serC* cells specifically engineered to autonomously biosynthesize nhpSer from the central metabolite phosphoenolpyruvate ^32^. It was reported that phosphonomethylalanine (nhpSer) effectively imitates phosphoserine in previous 14-3-3 binding studies ^32,34^. Likewise, we also produced the SARS-CoV-2 N protein containing only nhpSer205 imitating site-specifically phosphorylated protein at position 205. Given that this position within the RGT^205^SPAR motif can be occupied by a serine in natural SARS-CoV-2 variants (e.g., GenBank accession numbers, UDA99618.1, QWC76623.1, QRG25797.1, QQX30013.1, QQN91819.1) and in related orthologs, e.g. in Bat-CoV HKU3 N ^26^, instead of the threonine in SARS-CoV-2 N, the use of nhpS205 version instead of pT205 was fully justified.

All N proteins, and Clu1 and Clu3 mutants of 14-3-3σ used for crystallization, were purified by subtractive immobilized metal affinity and size-exclusion chromatography, where two IMAC steps were separated by 3C cleavage of the His_6_-tag ^36^. To remove large amounts of contaminating nucleic acids non-specifically bound to N, we used a continuous washing step (50 column volumes) by 3 M NaCl before elution with imidazole during IMAC1. Most proteins were obtained at amounts exceeding 10 mg per 1 liter of bacterial culture. While the site-specifically modified SARS-CoV-2 N 197TAG and 205TAG were obtained at rather low yields (∼1-2 mg per 1 liter of bacterial culture) and were contaminated by truncated side-products, the resulting proteins were modified as expected according to Phos-tag gels (10% polyacrylamide gels with 50 µM Mn^2+^ Phos-tag (Wako)). Despite SARS-CoV-2 N pS197 was purified in the presence of inhibitors of proteases (EDTA-free Pierce™ Protease Inhibitor Mini Tablets, cat. no. A32955) and phosphatases (orthovanadate, glycerophosphate and NaF), the product was considerably dephosphorylated. This prompted us to obtain also the nhpS197 and nhpS205 versions of this protein that resist dephosphorylation. SARS-CoV N was expressed as a GST-fusion and purified by a combination of a GSTtrap step 1, 3C proteolysis to cut off GST, a GSTtrap step 2 and heparin-affinity chromatography. All purified N preparations used in this study showed an *A*_260_/*A*_280_ ratio of 0.57-0.6 and were free from nucleic acids.

Untagged 14-3-3γ was expressed in *E*.*coli* BL21(DE3) cells by IPTG induction (final concentration of 1 mM) and purified by ammonium sulfate fractionation, anion-exchange and size-exclusion chromatography as described earlier ^26^. Expression of the MBP-14-3-3 constructs was the same, but these proteins were purified using a combination of MBPTrap column and size-exclusion chromatography ^28^. Protein concentrations were determined spectrophotometrically at 280 nm on a NP80 nanophotometer (Implen, Germany) using extinction coefficients listed in Supplementary Table 5.

### Kinetics of N dephosphorylation

To assess the effect of 14-3-3γ binding on the kinetics of N dephosphorylation, we used full-length SARS-CoV N phosphorylated during co-expression with PKA, because this N preparation provided the best contrast between the phosphorylated and unphosphorylated states on Phos-tag gels, facilitating densitometry. The effect of 14-3-3γ was also preserved for phosphorylated SARS-CoV-2 N, but the densitometric analysis was less reliable in this case. The reaction mixture contained 25 µM of phospho-N, 60 µM 14-3-3γ and 2 U of shrimp alkaline phosphatase in 20 mM Tris-HCl pH 8.0 containing 50 mM NaCl and 10 mM MgCl_2_, and dephosphorylation was followed by taking aliquots of the mixture at different time points, quenching them by acetone precipitation and analyzing on Phos-tag (10% polyacrylamide gels with 50 µM Mn^2+^ Phos-tag) and SDS-PAGE gels. The control reaction did not contain 14-3-3γ. Phos-tag runs were followed by GFP fluorescence on a control lane. Densitometry was done in Image Lab 6.1 (Bio-Rad). The experiment was done four times with different amounts of phosphatase, which yielded qualitatively similar results.

### Analytical size-exclusion chromatography

The interaction of N proteins and 14-3-3γ was analyzed using a Superdex 200 Increase 5/150 column (GE Healthcare) operated at a 0.45 ml/min flow rate on a Prostar 335 chromatographic system (Varian, Australia). When required, fractions were collected and analyzed by SDS-PAGE. In specific cases, a Superdex 200 Increase 10/300 column and a 0.8 ml/min flow rate was used. The buffer, used to equilibrate the columns, contained 20 mM Tris-HCl pH 7.6, 3 mM NaN_3_ and 200-300 mM NaCl. We noticed that free unphosphorylated SARS-CoV-2 N.S197L absorbed onto the column when equilibrated with <220 mM NaCl. Therefore, the reported experiments contained at least this much salt in the SEC buffer. Experiments were done at least three times with qualitatively very similar results.

### Multi-angle light scattering (MALS)

To determine absolute masses of MBP-14-3-3ζ, 14-3-3γ and its complex with phosphorylated SARS-CoV-2 N.S197L, we used SEC coupled to MALS. To do this, a chromatographic system was connected to the miniDAWN detector (Wyatt Technology) calibrated relative to the scattering from toluene. The concentration signal was obtained from SEC traces recorded at 280 nm and weight extinction coefficients 1.38 for MBP-14-3-3ζ, 1.13 for 14-3-3γ, and 1.02 for the 14-3-3γ/pN.S197L complex. The *M*_w_ distributions were calculated in Astra 8.0 software (Wyatt Technology) using dn/dc=0.185. Protein content in the peaks was additionally analyzed by SDS-PAGE.

### Fluorescence anisotropy

Fluorescence anisotropy (FA) was measured at 25 °C on a Clariostar plus microplate reader (BMG Labtech, Offenburg, Germany) using a set of bandpass filters (482±16 nm and 530±40 nm), FITC labeled phosphopeptides (100 nM, on a 20 mM HEPES-NaOH buffer, pH 7.5, 150 mM NaCl, 0.02% Tween 20) and 384-well plates (Black Nunc™ 384-Shallow Well Standard Height Polypropylene Storage Microplates, Thermofisher scientific, cat. no. 267460). A series of concentrations of MBP-14-3-3 proteins in a 20 mM HEPES-NaOH buffer, pH 7.5, 150 mM NaCl, 0.01% Tween 20 (33 μl each) was diluted 2-fold by a 100 nM solution of either FITC-labeled peptide to obtain 66 μl samples containing 50 nM peptide, MBP-14-3-3 (from 0 to 110 μM) and 0.01% Tween 20. After 5 min incubation at room temperature the fluorescence anisotropy readout did not change with time, indicating equilibrium. Then, each sample was split into three 20 μl aliquots, the plates were centrifuged for 1 min at 25 °C at 250 g to remove air bubbles, and the FP data were recorded. The triplicate measurements were converted into mean ± standard deviation values, which were used for the data presentation and fitting. To determine *K*_D_ values, the binding curves were approximated using the quadratic equation ^50^ in Origin 9.0 (OriginLab Corporation, Northampton, MA, USA). Selectivity coefficient reflecting the preferred 14-3-3 binding to the pT205 peptide than the pS197 peptide, which by definition is an equilibrium constant (*K*_eq_ = 1/*K*_D_) for the corresponding displacement reaction, was determined as the ratio of *K*_D_ for the pS197 peptide to *K*_D_ for the pT205 peptide. To determine the correlation between selectivity or -log*K*_D_ values for the 14-3-3/phosphopeptide pairs and average ΔΔ*G* values characterizing the affinity hierarchy reported earlier ^28^, we used the standard Pearson’s correlation coefficient *r* and coefficient of determination *R*^*2*^.

### Crystallization and X-ray data collection

Crystallization was performed using 14-3-3σ surface entropy reduction mutants Clu1 or Clu3 (storage buffer 20 mM Tris-HCl pH 7.6, 150 mM NaCl, 0.1 mM EDTA, 2 mM DTT, 2 mM MgCl_2_) mixed with either pS197 or pT205 phosphopeptide at a 1:5 protein:peptide molar ratio with the final protein concentration of 11.5 mg/ml. The best diffracting crystal for the 14-3-3σ Clu1/pT205 complex grew in condition H7 (0.2 M AmSO_4_, 100 mM BisTris pH 5.5, 25% PEG 3350) of JCSG+ screen (Molecular Dimensions, Sheffield, UK); the best diffracting crystal for the 14-3-3σ Clu3/pS197 complex grew in condition G2 (0.2 M NaBr, 100 mM BisTris propane pH 7.5, 20% PEG 3350) of PACT screen (Qiagen, Hilden, Germany). Crystals were mounted in nylon loops directly from crystallization drops and flash frozen in liquid nitrogen without additional cryoprotection. X-ray data (Supplementary Table 2) were collected at 100 K at the Diamond Light Source (Oxfordshire, UK) and processed using DIALS ^51^.

### Crystal structure determination and refinement

X-ray structures were determined by MolRep ^52^ using an unliganded 14-3-3σ dimer (PBD 5LU2) as a search model. Peptide residues were manually built into the difference electron density maps using Coot ^53^. Structures were refined by Buster 2.10.4 ^54^ imposing NCS restraints and TLS and all-atom individual isotropic B-factor restrained refinement. The refined structures were validated using the validation algorithm in Phenix 1.10 ^55^. Structural illustrations were prepared using PyMOL 2.20 (Schrodinger, Inc.).

### SEC coupled with small-angle X-ray scattering (SAXS)

SAXS data (*I*(*s*) versus *s*, where *s* = 4πsinθ/λ, 2θ is the scattering angle and λ = 0.96787 Å) from the mixture of phosphorylated SARS-CoV-2 N S197L mutant with a 1.7x molar excess of 14-3-3γ (45 µl, 10 mg/ml), loaded onto a Superdex 200 Increase 3.2/300 column (Cytiva) and eluted at a 0.075 ml/min flow rate, were measured at the BM29 beam line (ESRF, Grenoble, France) using a Pilatus 2M detector (data collection rate 0.5 frame/s; experiment session data DOI 10.15151/ESRF-ES-642726753). The running buffer contained 20 mM Tris-HCl, pH 7.6, and 150 mM NaCl. One thousand 1D SAXS frames acquired along the SEC profile were analyzed using CHROMIXS ^56^ to get a SAXS curve corresponding to the peak of the protein complex. This SAXS curve was further analyzed using ATSAS 2.8 program package ^42^. Because of the tailing peak of the complex, the peak of free 14-3-3γ became contaminated and was not analyzed. To perform structural modeling of the complex, we took available crystallographic data for the SARS-CoV-2 N NTD monomer (residues 48-175, PDB 7CDZ), CTD dimer (residues 254-364, PDB 6WZO) and also considered as rigid body the well-documented hydrophobic α1-helix within the interdomain linker (residues 220-232, PDB 7PKU) ^41^. In our 14-3-3σ dimer structure with two SARS-CoV-2 N pT205 peptides (residues 203-209), we replaced 14-3-3σ by the 14-3-3γ dimer (residues 1-235, PDB 6A5S). Considering rigid bodies above, we supplemented components of the assembly with missing N-terminal, C-terminal and linker residues absent from their atomistic structures (totaling ∼26% of all 1338 residues in the complex), and systematically changed relative positions of two NTD protomers, two α1 helices and CTD dimer relative to the 2:2 14-3-3γ/pT205 peptide complex using CORAL ^42^, to minimize the discrepancy between the calculated scattering profile and the experimental data. This fitting procedure showed high convergence (χ^2^ for all 20 models generated were close to 1). Alternative situations, represented by individual 14-3-3γ dimer or SARS-CoV-2 N dimer with supplemented flexible residues could not describe SAXS data for the complex (χ^2^>200 and 4.5, respectively).

## Acknowledgements

We thank Prof. Gilles Trave’s laboratory for providing the MBP-14-3-3 plasmids, Prof. Fulvio Reggiori for the SARS-CoV N plasmid, Dr. Olga Moroz for help with crystallization, Dorothy Hawkins for help with protein purification, Sam Hart, Anton Popov and Dmitry Soloviov for help with X-ray data collection and Dr. Gergo Gogl for advice on fluorescence anisotropy. LC-MS and MALS were carried out at the Shared-Access Equipment Centre “Industrial Biotechnology” of the Federal Research Center “Fundamentals of Biotechnology” of the Russian Academy of Sciences. The study was supported by the Ministry of Science and Higher education of the Russian Federation in the framework of the Agreement no. 075-15-2021-1354 (07.10.2021) and the Wellcome Trust (206377 award to A.A.A.). Expression and purification of 14-3-3 proteins were partially supported by the Russian Science Foundation (no. 19-74-10031). R.B.C. acknowledges support by the Medical Research Foundation at the Oregon Health Sciences University (USA) and the Collins Medical Trust, and also support by the GCE4All Biomedical Technology Development and Dissemination Center supported by National Institute of General Medical Science grant RM1-GM144227.

## Author Contributions

NNS and AAA conceived studies; PZ and RBC – prepared pSer and nhpSer incorporating cells; KVT, AAS, AAK, JLRS, NNS – expressed and purified proteins; KVT – performed fluorescence anisotropy experiments; JLRS – crystallized proteins; NNS – determined crystal structures and did SAXS modeling; KVT, RBC, AAA, NNS – analyzed data; NNS and KVT made illustrations; NNS wrote the paper with input from RBC and AAA.

## Data and materials availability

The refined models and structure factors have been deposited in the Protein Data Bank under the accession codes 7QIK and 7QIP. All materials are available from the corresponding authors upon reasonable request.

