## Supplementary Tables 1-5 Supplementary Figs 1-9 for "Human 14-3-3 proteins site-selectively bind the mutational hotspot region of SARS-CoV-2 nucleoprotein modulating its phosphoregulation"

**Supplementary information**

*Tugaeva et al.*

Contents:

**Supplementary Tables**

**Supplementary Table 1.** Apparent *K*_D_ values and selectivity coefficients for the interaction of seven 14-3-3 isoforms with two SARS-CoV-2 N phosphopeptides

**Supplementary Table 2.** X-ray data collection and refinement statistics.

**Supplementary Table 3.** Structural parameters of the 14-3-3γ/SARS-CoV-2 N complex in solution based on SEC-SAXS data.

**Supplementary Table 4**. Monoisotopic masses of the synthetic peptides and their FITC-labeled derivatives used in the study.

**Supplementary Table 5**. Parameters of protein constructs used in this study.

**Supplementary Figures**

**Supplementary Fig. 1**. SARS-CoV N interacts with human 14-3-3γ in a phosphorylation-dependent manner.

**Supplementary Fig. 2**. The S197L mutant of SARS-CoV-2 N interacts with human 14-3-3γ in a phosphorylation-dependent manner.

**Supplementary Fig. 3**. SEC-MALS analysis of the MBP-14-3-3ζ fusion.

**Supplementary Fig. 4**. SARS-CoV-2 N phosphopeptides interact with the common peptide-binding 14-3-3 grooves.

**Supplementary Fig. 5**. Titration of SARS-CoV-2 N phosphopeptides by the seven human 14-3-3 isoforms monitored by fluorescence anisotropy.

**Supplementary Fig. 6.** The six core residues in the two SARS-CoV-2 N phosphopeptides form similar contacts with 14-3-3σ.

**Supplementary Fig. 7.** The unique salt bridge-forming glutamate residue is highly conserved in the human 14-3-3 protein family.

**Supplementary Fig. 8**. MS-based validation of the peptides and their labeled derivatives.

**Supplementary Fig. 9.** Sequencing results for the S197L and S197TAG mutants of SARS-CoV-2 N.

**Supplementary Tables**

**Supplementary Table 1.** Apparent *K*_D_ values and selectivity coefficients for the interaction of seven 14-3-3 isoforms with two SARS-CoV-2 N phosphopeptides as determined by fluorescence anisotropy.

| 14-3-3 isoform | *K*_D_ ±SD, µM | | Selectivity* |
| --- | --- | --- | --- |
|  | pT205 | pS197 |  |
| gamma | 10.7 ±0.3 | 71.6 ±1.0 | 6.7 |
| eta | 13.2 ±0.3 | 72.0 ±1.2 | 5.5 |
| beta | 14.4 ±0.3 | 69.9 ±1.5 | 4.8 |
| epsilon | 17.7 ±0.4 | 83.7 ±1.4 | 4.7 |
| zeta | 19.2 ±0.4 | 81.4 ±1.4 | 4.2 |
| tau | 20.8 ±0.4 | 81.9 ±1.5 | 3.9 |
| sigma | 32.9 ±0.8 | 98.6 ±1.9 | 3.0 |

* selectivity coefficients characterizing the preferred binding of the pT205 peptide over the pS197 peptide were calculated for the given isoform as the ratio of the *K*_D_ for the pS197 to the *K*_D_ for the pT205 peptide.

**Supplementary Table 2.** X-ray data collection and refinement statistics.

|  | 14-3-3σ/pS197 structure | 14-3-3σ/pT205 structure |
| --- | --- | --- |
| *Data collection* | | |
| Detector | Dectris EIGER2 Si 16M | Dectris EIGER2 XE 16M |
| Wavelength, Å | 0.9795 | 0.9763 |
| Resolution range, Å | 57.60-2.01 (2.12 - 2.01)* | 70.45 - 2.65 (2.70 - 2.65)* |
| Space group | P2_1_2_1_2_1_ | P4_1_ |
| Cell parameters : a, b, c (Å) | 47.4, 111.3, 115.2 | 99.7, 99.7, 58.6 |
| Total reflections | 418,896 (19,731)** | 200,790 (10,676) |
| Unique reflections | 39,958 (4740) | 16,912 (818) |
| Multiplicity | 10.5 (4.2) | 11.9 (13.1) |
| Completeness (%) | 96.6 (80.8) | 100 (100) |
| Mean I/σ(I) | 11.2 (1.3) | 4.8 (0.9) |
| R-meas | 10.5 (97.6) | 36.4 (225.8) |
| CC_1/2_ | 0.998 (0.767) | 0.989 (0.353) |
| *Refinement* | | |
| R-work | 0.192 *** | 0.221 *** |
| R-free | 0.217 | 0.251 |
| Number of non-H atoms | 3918 | 3755 |
| Protein atoms | 3580 | 3607 |
| Heteroatoms | 68 | 25 |
| Solvent atoms | 338 | 115 |
| Protein residues | 459 | 471 |
| RMS(bonds), Å | 0.014 | 0.014 |
| RMS(angles), ° | 1.4 | 1.4 |
| Ramachandran favored (%) | 98.9 | 98.0 |
| Ramachandran outliers (%) | 0 | 0.2 |
| Rotamer outliers (%) | 4.0 | 4.3 |
| C beta outliers | 0 | 0 |
| Clashscore | 0.54 | 0.83 |
| MolProbity overall score | 1.14 | 1.24 |
| Average B-factor | 47.6 | 70.3 |
| PDB ID | 7QIK | 7QIP |

* Statistics for the outer shell are in parentheses.** Data processing was using DIALS ^1^. *** Data are from validation using Phenix ^2^.

**Supplementary Table 3.** Structural parameters of the 14-3-3γ/SARS-CoV-2 N complex in solution based on SEC-SAXS data.

|  | 14-3-3γ/phospho-SARS-CoV-2 N S197L |
| --- | --- |
| Protein concentration loaded, mg ml^-1^ | 10 |
| **Guinier analysis** | |
| *I*(0) | 42.44 ±0.17 |
| *R*_g_ (nm) | 5.44 ± 0.03 |
| s*R*_g_ range | 0.7 < *sR*_g_ < 1.3 |
| *p*(r) analysis |  |
| *R*_g_ (nm) | 5.69 ± 0.04 |
| *R*_max_ (nm) | 20.0 |
| *s* range (nm^−1^) | 0.15–2.50 |
| Kratky plot | bell-shaped, with shoulder and significant flexibility |
| **Volume, shape and molecular weight (*M*_W_) analysis** | |
| Porod volume, nm^3^ | 320.4 |
| *M*_W_ calculated from amino acid sequence, kDa | 148.3 (for 2:2 complex) |
| *M*w from SEC-MALS, kDa (*M*_W_ ratio) ^a^ | 150.8 (1.02) |
| *M*_W_ from Porod volume, kDa (*M*_W_ ratio) | 188.5 (1.27) |
| *M*_W_ from SAXSMoW, kDa (*M*_W_ ratio) | 144.3 (0.97) |
| *M*_W_ from *V*_c_, kDa, (*M*_W_ ratio) | 153.3 (1.03) |
| *M*_W_ from Bayesian inference, kDa, (*M*_W_ ratio) | 169.6 (1.14) |
| CORAL | 20 calculations |
| *χ*^2^ range (all models) | 0.94-1.05 |
| *s* range for fitting (nm^−1^) | 0.13-2.58 |
| **CRYSOL (50 harmonics, 256 points, constant enabled) ^b^** | |
| *s* range for model fitting (nm^−1^) | 0.13–5.01 |
| *χ*^2^ (the best model) | 1.05 |
| model *R*_g_ (nm) | 5.80 |

^a^The experimental *M*_W_ ratio relative to the *M*_W_ calculated for a 2:2 complex. ^b^CRYSOL ^3^ fits to the SAXS data for the whole range of scattering vectors. All analysis was done using programs included in the ATSAS 2.8 software package ^4^.

**Supplementary Table 4**. Monoisotopic masses of the synthetic peptides and their FITC-labeled derivatives used in the study.

| peptide | | *M*_W_ (expected), Da | | *M*_W_ (according to LC-MS), Da | |
| --- | --- | --- | --- | --- | --- |
|  | sequence | unlabeled | labeled by FITC | unlabeled | labeled by FITC |
| pS197 | WSSRN**pS**TPGSS | 1244.47 | 1633.51 | 1244.49 | 1633.51 |
| pT205 | WSSRG**pT**SPARM | 1314.54 | 1703.58 | 1314.58 | 1703.59 |
| pB6 | WLRRA**pS**APLPGLK | 1543.83 | 1932.86 | 1543.84 | 1932.87 |

**Supplementary Table 5**. Parameters of protein constructs used in this study.

| Protein | *M*_W_, Da | pI | Molar ext. coef.  M^-1^ cm^-1^ | Ext. coef.  (mg/mL)^-1^ cm^-1^ |
| --- | --- | --- | --- | --- |
| MBP-14-3-3ε | 69505.8 | 4.8 | 95230 | 1.4 |
| MBP-14-3-3γ | 68634.5 | 4.9 | 98210 | 1.4 |
| MBP-14-3-3ζ | 68077.0 | 4.9 | 93740 | 1.4 |
| MBP-14-3-3τ | 68298.4 | 4.9 | 93740 | 1.4 |
| MBP-14-3-3η | 68752.9 | 4.9 | 95230 | 1.4 |
| MBP-14-3-3σ | 68308.2 | 4.8 | 92250 | 1.4 |
| MBP-14-3-3β | 68616.5 | 4.9 | 93740 | 1.4 |
| 14-3-3γ | 28302.6 | 4.8 | 31860 | 1.1 |
| 14-3-3σ (1-231 aa) (^159^KKE^161^-^159^AAA^161^, Clu1) | 26225.5 | 4.8 | 25900 | 1.0 |
| 14-3-3σ (1-231 aa) (^75^EEK^77^-^75^AAA^77^, Clu3) | 26224.6 | 4.9 | 25900 | 1.0 |
| SARS-CoV N (without tag) | 46607.9 | 10.1 | 43890 | 0.9 |
| SARS-CoV-2 N (without tag) | 45851.0 | 10.1 | 43890 | 0.9 |
| SARS-CoV-2 N pSer197 (without tag) | 46492.5 | - | 43890 | 0.9 |
| SARS-CoV-2 N nhpSer197 (without tag) | 46490.6 | - | 43890 | 0.9 |
| SARS-CoV-2 N nhpSer205 (without tag) | 46476.6 | - | 43890 | 0.9 |
| SARS-CoV-2 N S197L (without tag) | 45877.0 | 10.1 | 43890 | 1.0 |

**Supplementary Figures**

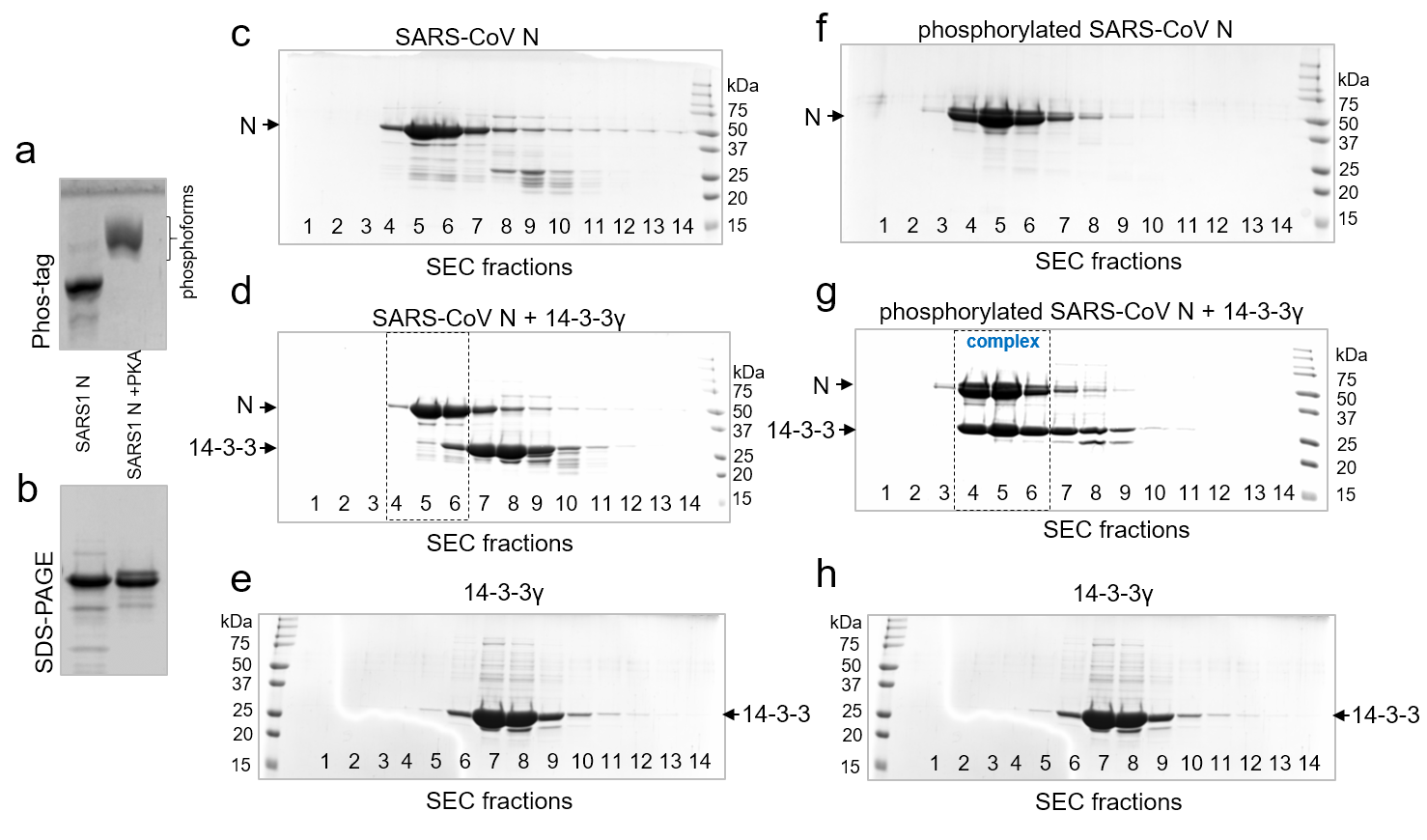

**Supplementary Fig. 1**. SARS-CoV N interacts with human 14-3-3γ in a phosphorylation-dependent manner. a,b. Phosphorylation status of SARS-CoV N co-expressed with PKA in *E.coli* as analyzed by Phos-tag and SDS-PAGE. c-e. The interaction of unphosphorylated SARS-CoV N with human 14-3-3γ analyzed by SEC with SDS-PAGE of all fractions along the elution profiles. f-h. The interaction of PKA-phosphorylated SARS-CoV N with human 14-3-3γ analyzed by SEC with SDS-PAGE of all fractions along the elution profiles. c and f correspond to SEC profiles of N or pN. d and g correspond to SEC profiles of 14-3-3 mixtures with either N or pN. e and h correspond to a SEC profile of individual 14-3-3γ. All SEC runs were performed at a 14-3-3 excess and at 220 mM NaCl in the samples and running buffer. Molecular mass markers are indicated in kDa. Positions of proteins are indicated by arrows. Note that only phosphorylated SARS-CoV N forms a complex with 14-3-3γ eluting in fractions 4-6 outlined by dashed rectangles.

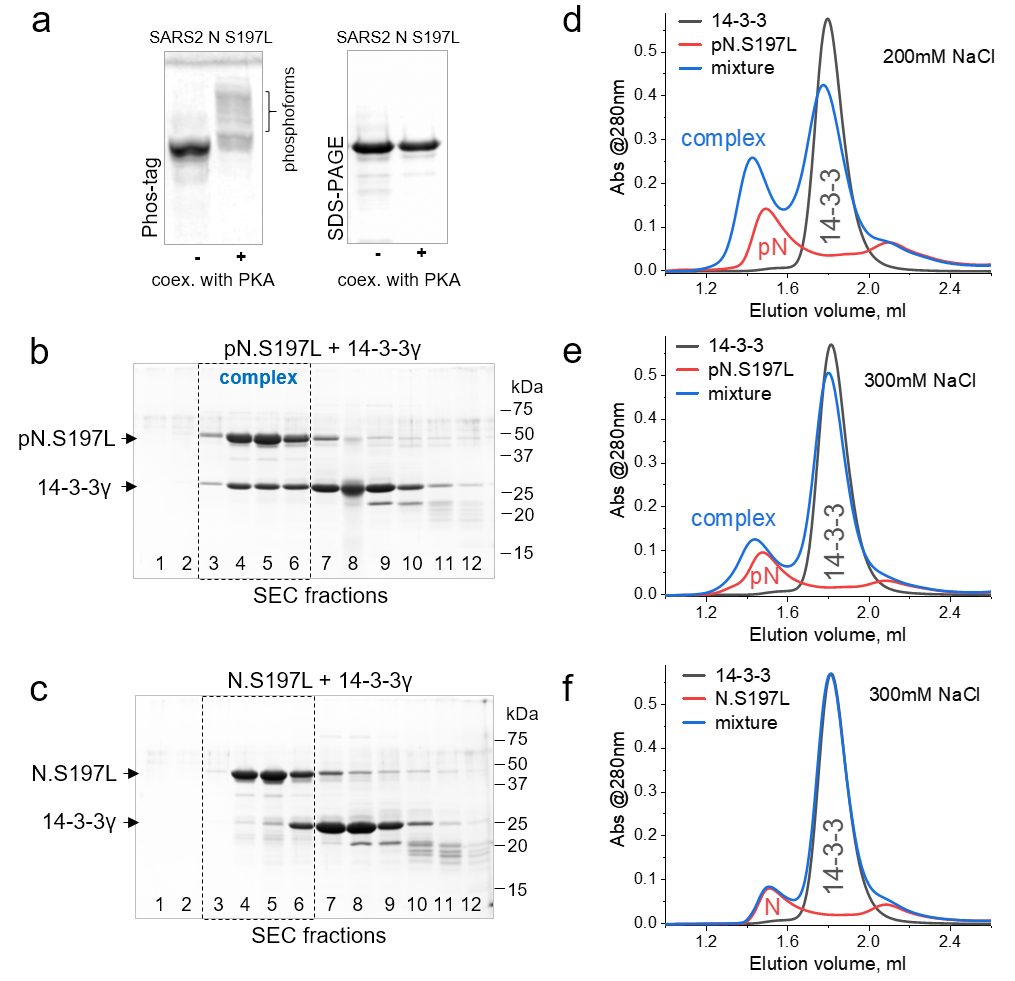

**Supplementary Fig. 2**. The S197L mutant of SARS-CoV-2 N interacts with human 14-3-3γ in a phosphorylation-dependent manner. a. Phosphorylation of SARS-CoV-2 N variant analyzed by Phos-tag (left) and SDS-PAGE (right). b,c. The interaction of human 14-3-3γ with PKA-phosphorylated (b) or unphosphorylated (c) SARS-CoV-2 N.S197L mutant analyzed by SEC and SDS-PAGE of the eluted fractions. 14-3-3γ was pre-incubated in excess with N in 220 mM NaCl, and SEC was run in the same buffer. Molecular mass markers are indicated in kDa on the right. Positions of proteins are indicated by arrows on the left. Note that fractions 3-6 (dashed rectangles) contained the 14-3-3:N complex only in the case of pN.S197L. d-f. Effect of salt on the interaction. SEC profiles of 14-3-3γ, phosphorylated N.S197L or their mixture, obtained at 200 mM NaCl (d) or 300 mM NaCl (e) in the premixed samples and running buffer. Note that the complex is detectable even at 300 mM NaCl. f. 300 mM NaCl in the premixed samples and SEC buffer fully blocks nonspecific interaction of 14-3-3γ with unphosphorylated N.S197L.

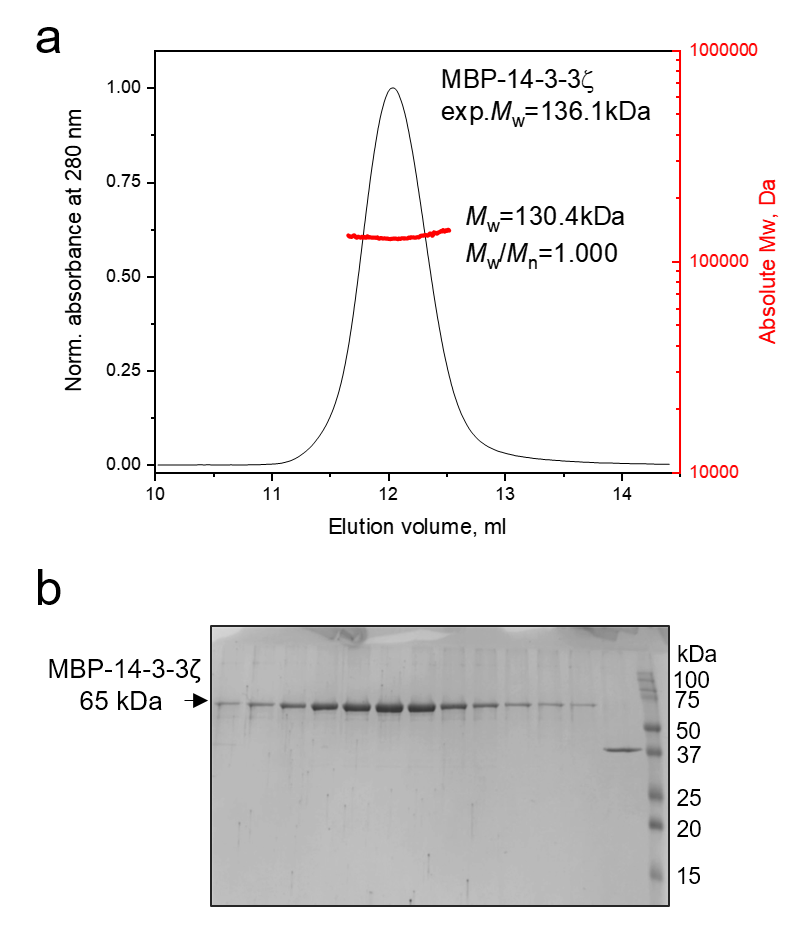

**Supplementary Fig. 3**. SEC-MALS analysis of the MBP-14-3-3ζ fusion. a. SEC-MALS profile from the Superdex 200 Increase 10/300 column followed by absorbance at 280 nm showing the *M*_w_ distribution across the protein peak as calculated from MALS data. Polydispersity index (*M*_w_/*M*_n_) and the expected *M*_w_ for the protein dimer are indicated. Flow rate was 0.8 ml/min. b. SDS-PAGE analysis of the fractions collected from the column. Mass markers are indicated in kDa. Protein position and its apparent mass are marked by an arrow.

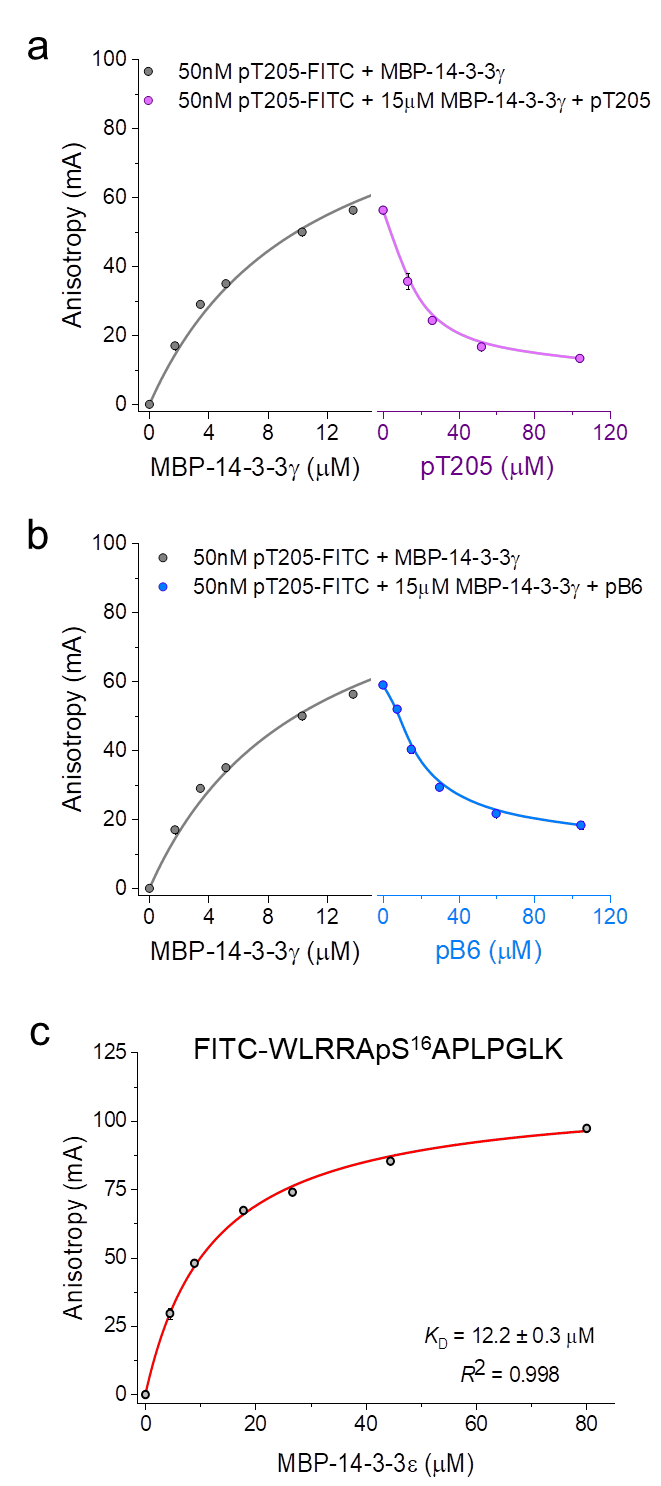

**Supplementary Fig. 4**. SARS-CoV-2 N phosphopeptides interact with the common peptide-binding 14-3-3 grooves. The FITC-labeled SARS-CoV-2 N pT205 phosphopeptide is outcompeted from its complex with human 14-3-3γ either by the same unlabeled phosphopeptide (a) or by the control unlabeled pB6 peptide (b) as monitored by fluorescence anisotropy. c. Titration of the FITC-labeled pB6 phosphopeptide by MBP-14-3-3ε, with the corresponding *K*_D_. In a-c, background anisotropy was subtracted from all values. The data are shown as mean ± standard deviation (*n*=3).

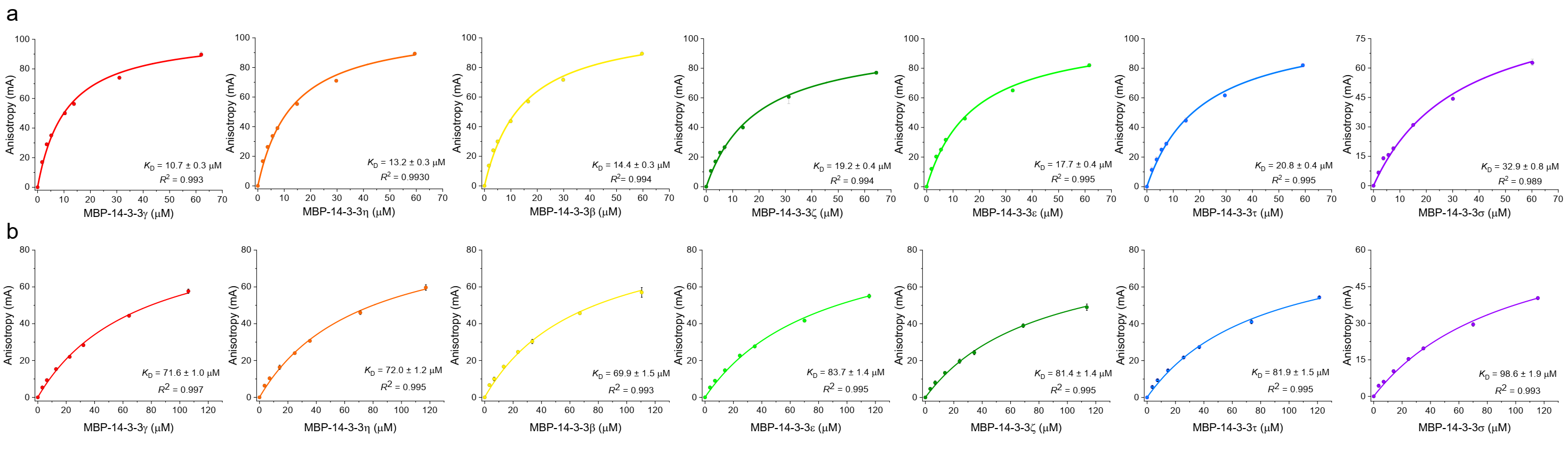

**Supplementary Fig. 5**. Titration of SARS-CoV-2 N phosphopeptides by the seven human 14-3-3 isoforms monitored by fluorescence anisotropy. a. Binding curves for the pT205 peptide. b. Binding curves for the pS197 peptide. Fitting of the curves was done in Origin 9.0 by standard quadratic binding equation ^56^. Apparent dissociation constants (*K*_D_) at equilibrium are shown on the panels along with the coefficient of determination *R*^2^. The data are shown as mean ± standard deviation (*n*=3). Fluorescence anisotropy data throughout titrations were corrected by subtracting the background anisotropy values.

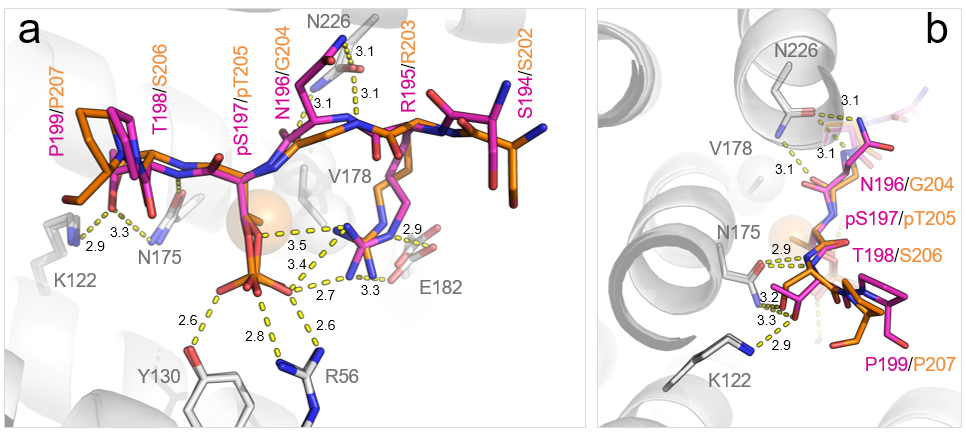

**Supplementary Fig. 6.** The six core residues in the two SARS-CoV-2 N phosphopeptides form similar contacts with 14-3-3σ. a, b. Superimposition of the two crystal structures showing the similarity of the conformation of both peptides and molecular contacts formed, looking at two different angles. H-bonds are shown by yellow dashed lines, hydrophobic interaction between the methyl group unique to pT205 and the side chain of Val178 of 14-3-3σ is shown by large semi-transparent spheres, distances characterizing key polar contacts are indicated in Å. Note that the hydrophobic contact made by the methyl group of pT205 is somewhat compensated in the pS197 peptide by an additional H-bond from the pS197 peptide’s Asn196 side chain and the side chain of Asn226 of 14-3-3σ, making the interaction of the core part of the peptides nearly equivalent in terms of the amount of chemical contacts.

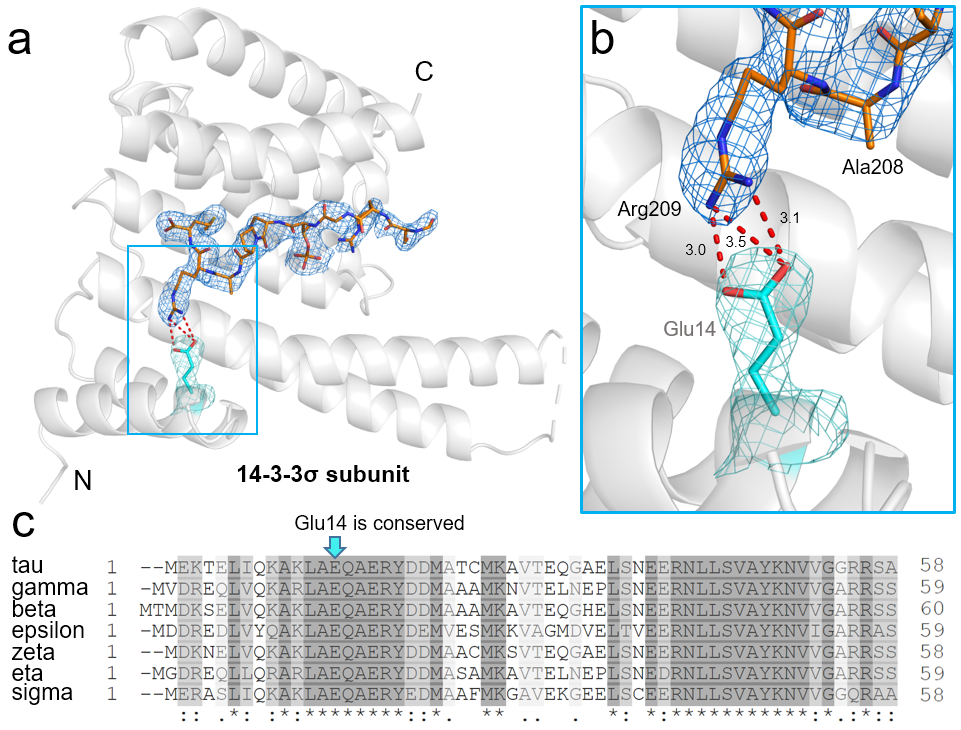

**Supplementary Fig. 7.** The unique salt bridge-forming glutamate residue is highly conserved in the human 14-3-3 protein family. a. Crystal structure of 14-3-3σ subunit (grey cartoon) complexed with the pT205 phosphopeptide of SARS-CoV-2 N (orange sticks) showing the location of the unique salt bridge (red dashed lines). 2Fo-Fc electron density map for the analyzed region is contoured at 1σ. b. Close-up of the salt bridge between the side chains of Arg209 of the SARS-CoV-2 N peptide and Glu14 of 14-3-3σ showing characteristic distances in Å. 2Fo-Fc electron density map for the analyzed region is contoured at 1σ. c. Local multiple sequence alignment of the human 14-3-3 isoforms in their N-terminal region, showing the absolute conservation of the glutamate partaking in the salt bridge to Arg209 of SARS-CoV-2 N. To the best of our knowledge, this glutamate is conserved in all 14-3-3 proteins. This implies that the ability to stabilize the binding of the pT205 phosphopeptide of SARS-CoV-2 N is shared by all isoforms with no exception.

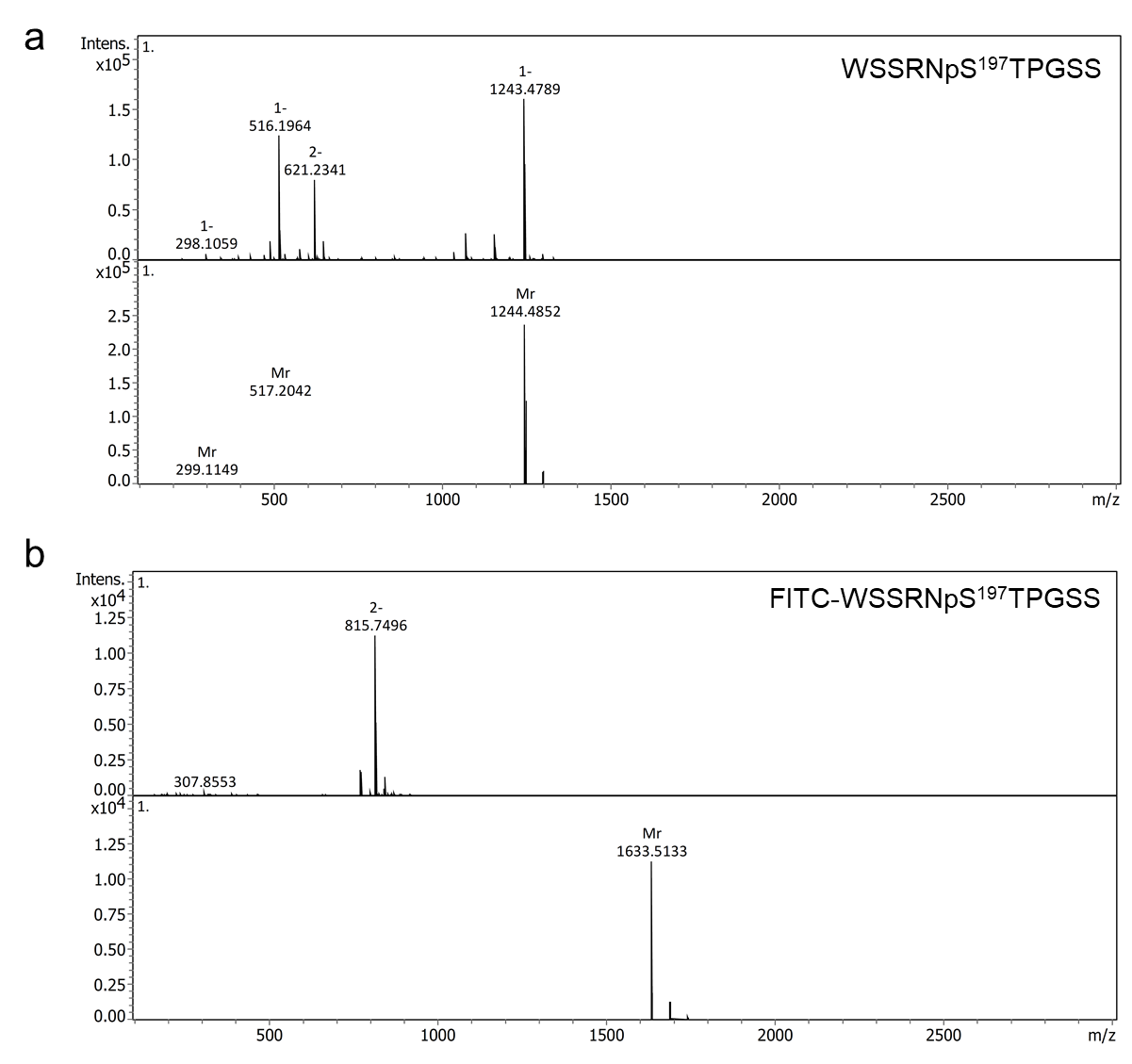

**Supplementary Fig. 8**. MS-based validation of the peptides and their labeled derivatives. LC-MS spectra for the pS197 peptide (a) and its FITC-labeled derivative (b).

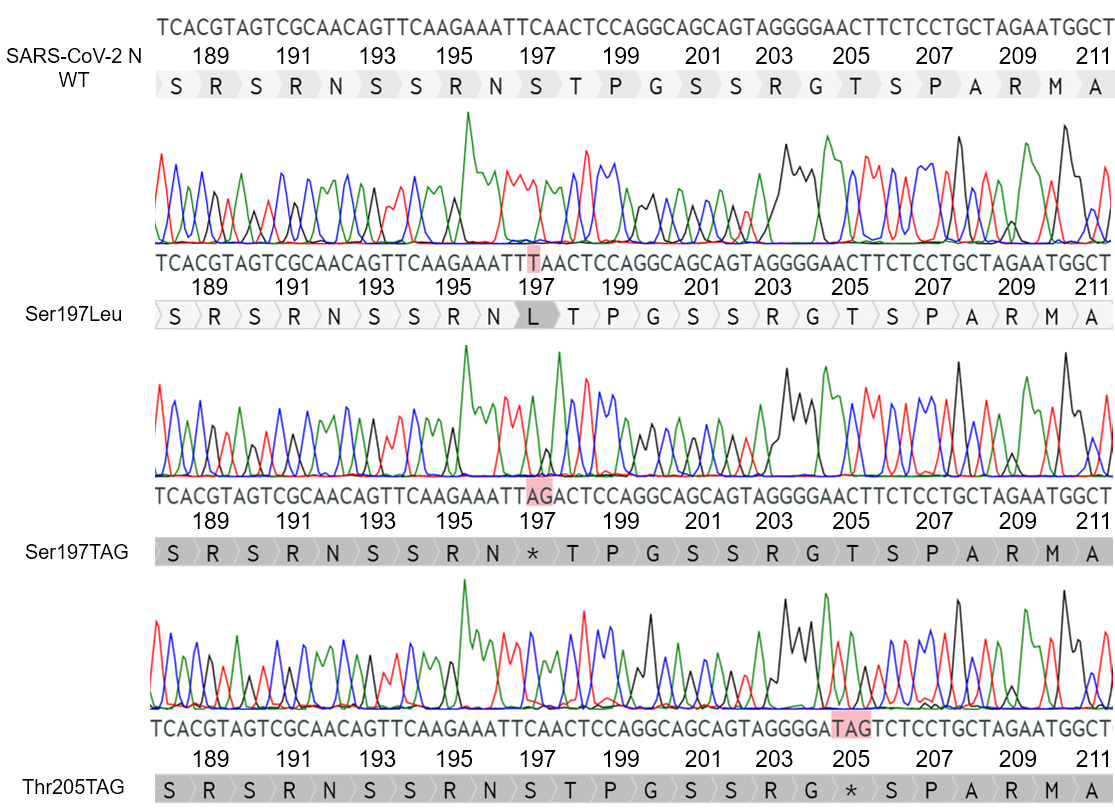

**Supplementary Fig. 9.** Sequencing results for the S197L, S197TAG and T205TAG mutants of SARS-CoV-2 N. Chromatogram peaks color: A – green, T – red, C – blue, G – black. The introduced replacements are highlighted by red. On top, the nucleotide and amino acid sequences for the wild-type protein are shown for reference.
